# Recruitment of apolipoprotein E facilitates Herpes simplex virus 1 attachment, entry, and release

**DOI:** 10.1101/2023.02.10.526562

**Authors:** Lifeng Liu, Konrad Thorsteinsson, Fouzia Bano, Sanduni Wasana Jayaweera, Damien Avinens, Dario Conca, Hudson Pace, Hugo Lövheim, Anders Olofsson, Marta Bally

**Author notes:** Correspondence: Lifeng Liu,; Marta Bally,. Department of Clinical Microbiology, Umeå University, Sweden. These authors contribute equally to this work (Konrad Thorsteinsson, Fouzia Bano).

## Abstract

Over two decades, epidemiological studies have revealed that interactions between human polymorphic apolipoprotein 4 (ApoE, isoform 4) and herpes simplex virus type 1 (HSV1) associate with higher risk of Alzheimer’s disease, a serious and increasing issue among elder populations worldwide. Nevertheless, little is known about the mechanisms behind ApoE-HSV1 interactions at molecular levels. Here, we investigate the effects of ApoE on the HSV1 infectious life cycle in *in vitro* cell experiments. Analysis of HSV1 growth curves shows that HSV1 production is promoted in presence of any of the three ApoE isoforms, with ApoE 3 or 4 demonstrating more proviral effects than ApoE 2. Quantification by qPCR reveals that the presence of ApoE 2, 3, or 4 leads to an increase of HSV1 extracellular release but unchanged levels of viral genome copies within cells or on the cell surface, indicating that virus replication, assembly, or transport to cell membrane are not affected. Further test of virus release directly demonstrates that HSV1 detachment from the cell surface is promoted by ApoE. Subsequent results reveal that ApoE is both present in purified HSV1 particles produced in ApoE-expressing cells after ultra-centrifugation and able to incorporate into HSV1 particles after purification, suggesting that harbouring ApoE may play a key role in the pro-viral effect of ApoE. Along these lines, we tested the infectious behaviour of ApoE-coated viruses and observed faster attachment kinetics and higher entry efficiencies of ApoE decorated HSV1. Our hypothesis that the association with ApoE leads to modified interactions of the virus with the cell membrane during entry and egress, was further validated in biophysical experiments. In such experiments, HSV1-membrane interaction kinetics and apparent affinity between HSV1 and native supported lipid bilayers (a plasma membrane mimic) were quantified using total internal reflection microscopy. HSV1 particles decorated with ApoE demonstrate both higher association (*k*_on_) and dissociation rate constants (*k*_off_), as well as less irreversible binding to the membrane, which is in line with the biological experiments. Overall, our results provide new insights into the roles of ApoE during HSV1 infections, which is worth to be considered when studying their involvement during Alzheimer’s disease development.

## Introduction

A possible involvement of herpes simplex virus 1 (HSV1) in the development of Alzheimer’s disease (AD), a neurodegenerative disease with increasing threats among elder populations, was first suggested in the 80s (Ball, 1982); in the later years, epidemiological studies have strengthened this hypothesis. One key finding of population-based cohort studies is the likely interaction effect of HSV1 and the apolipoprotein E allele 4 (E4) *APOE4* on AD, where these factors together associate with an increased risk of developing AD (Itzhaki et al., 2001; Itzhaki et al., 1997; Lin et al., 1996; Linard et al., 2020; Lindman et al., 2019; Lövheim et al., 2019; Steel & Eslick, 2015; Strandberg et al., 2005). Given that *APOE4* is such an established and strong AD risk factor, this interaction in turns provides strong support for a role of HSV1 infection in AD processes. Along these lines, a few studies have also linked *APOE* genotype to cold sore frequency or anti-HSV IgM antibody presence, indicative of an effect on the immunological control over HSV1(LIN et al., 1995; Lindman et al., 2022; Pandey et al., 2020). Still, considering the potential importance of the interaction between *APOE* genotype and HSV1 for AD risk, surprisingly little is known about how human apolipoprotein E (ApoE) and HSV1 interact mechanistically during virus infection.

Human apolipoprotein E (ApoE) is a polymorphic and multi-functional protein, 299 amino acids in length and containing two structural domains identified as the N-terminal (amino acids 1 – 191) and C-terminal domains (amino acids ∼ 206 – 299), connected by a linker. Functional regions located in the two domains include the receptor binding and heparan sulfate (HS) binding regions in the N-terminal domain (Libeu et al., 2001), as well as the lipid binding and self-association regions in the C-terminal domain (reviewed in (Tudorache et al., 2017)). The *ApoE* gene is located on human chromatin 19 (Myklebost & Rogne, 1988) and produces three major alleles, that encode three corresponding protein products termed ApoE 2, ApoE 3, and ApoE 4. These three ApoE isoforms differ at only two amino acid residues: 112 and 158; ApoE 3 has a cysteine at residue 112 and an arginine at 158; while at both sites ApoE 2 has cysteines and ApoE 4 has arginines. These single amino acid substitutions lead to important changes in the ApoE structure which in turn results in differences in the characteristics of the protein’s receptor, lipid and HS binding abilities (reviewed in (Mahley & Rall Jr., 2000)). For example, ApoE 2 shows significantly lower binding affinity than ApoE 3 or 4 to the low density lipoprotein (LDL) receptor on cultured cells, which is thought to be a result of the missing arginine at 158 (Dong et al., 1996). The arginine 112 in ApoE 4 enables unique domain interactions, accounting for its preference to very low density lipoproteins (VLDL), in comparison to binding to high density lipoprotein by ApoE 2 or ApoE 3 (Dong & Weisgraber, 1996). In addition, a biomolecular interaction study using surface plasmon resonance has observed that ApoE 4 shows significant higher binding than ApoE 3 or ApoE 2 to the glycosaminoglycans (GAG) heparan sulfate and dermatan sulfate found on the cell surface, which is postulated to be a result of electrostatic interactions with the N-terminal domain (Yamauchi et al., 2008). The role of ApoE in biological processes has been investigated beyond its conventional function in cholesterol transport; the protein has been shown to play roles in immune regulation, in virus infection and in the development of cardiovascular and neurological diseases (summarized in (Tudorache et al., 2017)). For decades, carrying the allele *ApoE4* has been seen as the most important risk factor for the development of sporadic Alzheimer’s disease, (see review (Kim et al., 2009)). Nevertheless, mechanistic insights on why ApoE 4 increases AD susceptibility are widely lacking, and the protein alone is not a strong risk factor for AD development, suggesting the involvement of other factors, such as an infection history with some human neurotropic pathogens. Many studies have suggested that herpesvirus infection, and in particular infection with HSV1, could be one of the important pathogenic factors for the AD development (Wainberg et al., 2021).

HSV is a human neurotropic virus member of the alpha Herpesviridae subfamily, which exists as two major serotypes HSV1 and HSV2. The prevalence of infection is estimated to be as high as 67% for HSV1 and 13% for HSV2 among people under age 50 (“Herpes simplex virus”. World Health Organization. 31 January 2017). One of the biological features of HSV infection is that the virus can produce lytic or latent infections, depending on the target cell types. HSV1 establishes a lifelong latency in neurons specifically but undergoes lytic cycles in other cell types after reactivation (Brown, 2017). The HSV virion is composed of three structural parts: an icosahedral capsid enclosing a linear, double-stranded DNA viral genome; a layer of proteins called tegument surrounding the capsid shell, and a glycoprotein-containing lipid membrane envelope as the outermost layer, wrapping both the tegument and nucleocapsid. The HSV DNA genome, approximately 152 kilo base pairs in length, encodes about 80 viral proteins, among which at least 12 of them are glycoproteins displayed on the envelope. Consistent with the large variety of glycoproteins found in the virion, HSV1 entry is a complex and versatile process which depends on the infected tissue and on the infection circumstances. Prior to entry, virus attachment is mediated by the interactions between virus glycoprotein B and C and GAGs on the cell surface, HS (Shieh et al., 1992) or chondroitin sulfate proteoglycans (Banfield et al., 1995). Recent studies have characterized interactions between HSV glycoprotein B or C and various GAGs (Altgärde et al., 2015; Delguste et al., 2019; Peerboom et al., 2017; Trybala et al., 2000) and revealed the interactions are important not only for virus attachment, but also for virus particle transport to entry receptors on cell surface (Abidine et al., 2022). These studies also highlight that the characteristics of such interactions need to be tightly regulated, not only to ensure efficient virus recruitment during entry but also efficient virus release upon egress. Glycoprotein o-glycosylation (Altgärde et al., 2015; Olofsson et al., 2023; Trybala et al., 2021) but also increased heparinase expression upon infection (Hadigal et al., 2020; Hopkins et al., 2018b) have been proposed as important factors contributing to a balanced GAG-virus interaction at the cell surface.

Recent studies have revealed that ApoE plays important roles during infections with different viruses (Tudorache et al., 2017) and is involved in the corresponding pathogenesis and diseases (Kuhlmann et al., 2010). ApoE expression is upregulated in HIV-infected macrophages but inhibits HIV infection when overexpressed in the 293T cell line (Siddiqui et al., 2018). In contrast, hepatotropic virus hepatitis B (HBV) and hepatitis C (HCV) viruses require ApoE for efficient infection and virus production, and ApoE has also been co-purified together with HBV and HCV particles, suggesting incorporation of the protein into virus particles (Bankwitz et al., 2017; Chang et al., 2007; Jiang & Luo, 2009; Qiao & Luo, 2019). Mechanistic studies suggest that virus attachment to heparan sulfate proteoglycan (HSPG) on the cell surface is facilitated by ApoE for both HBV and HCV (Jiang et al., 2013; Li & Luo, 2021) and that virus assembly or egress of HCV also benefits from ApoE (Benga et al., 2010; Cun et al., 2010; Lee et al., 2014). However, none of these effects are isoform dependent. Isoform-dependent effects have been observed in some HSV1 studies. Transgenic mice studies show that in comparison to ApoE 3, ApoE 4 leads to more brain access (Burgos et al., 2006) and latency to HSV1 (Bhattacharjee et al., 2006; Miller & Federoff, 2008), as well as to more severe pathogenesis (Bhattacharjee et al., 2008). Moreover, ApoE derived peptides were found to inhibit HSV1 infection (Bhattacharjee et al., 2008; Bhattacharjee et al., 2009; Dobson et al., 2006), likely by targeting HSPGs, a shared cell-surface receptor for both ApoE and HSV1 (Saito et al., 2003; Shieh et al., 1992). On the other hand, proviral effects of ApoE have been suggested in ApoE knockout mice, where HSV1 infection and transmission is decreased in different organs, such as the spinal cord and brain (Burgos et al., 2002). In spite of pathogenesis studies in animal models, and although similar studies have been done with other viruses, such as HIV, HBV, or HCV, little is known about ApoE effects on HSV1 growth at molecular levels. Here we investigate HSV1 infection on cultured cell lines in the presence of different ApoE isoforms. Our results reveal that association of ApoE with HSV1 particles promotes infection by facilitating both virus attachment to and release from the cell surface.

## Results

### HSV1 growth is promoted by the presence of ApoE

To study how the presence of ApoE affects HSV1 infection and explore potential differences between the allelic isoforms of the protein, we tested HSV1 growth in the presence of various concentrations of ApoE in both the neuronal blastoma cell line SH-SY5Y, and an epithelial cell line commonly used for HSV1 studies, green monkey kidney cells (GMK). In both SH-SY5Y and GMK cells, ApoE expression levels were negligible as verified by western blot (Fig. S1). This is in line with the notion that ApoE is mainly secreted by the liver or by astrocytes in the brain (Elshourbagy et al., 1984). HSV1 growth was thus investigated by adding ectopically expressed and purified ApoE solutions to cells. In such experiments, solutions of isoform 2, 3 or 4 with designated concentrations were prepared in cell culture mediums and then added to cells after 1h inoculation of HSV1 at multiplicity of infection (MOI) 0.1. At 24 hours post infection (hpi), viruses released in the medium were harvested and titrated by plaque assays. The results show that HSV1 growth was increased with 5 µM ApoE 2, 3 or 4 both on SH-SY5Y and GMK, but not at lower concentrations of any of the apoE isoforms (Fig. 1A). The promotion of HSV1 growth by ApoE is also isoform dependent, with ApoE 3 or 4 demonstrating higher effect than ApoE 2. There was no effect of the diluent on HSV1 growth as compared to that of the medium group, confirming that the increase of virus amounts in the supernatant resulted from the presence of ApoE. Concomitantly, ApoE uptake at different concentrations was evaluated by immunofluorescence staining (Fig. S2a), revealing that at least 2.5 µM is needed to guarantee an effective uptake of the protein by most cells, which is highly correlated to the promoted HSV1 growth at these two concentrations (Fig. S2b). HSV1 growth was also tested on SH-SY5Y cells where apoE was present prior to and kept during the period of virus infection except for the 1 h inoculation. Promotion of HSV1 growth was observed with 5 µM ApoE 2, 3 or 4 to a similar extent as that in Fig. 1A, but not under other conditions (Fig. S3a), suggesting that apoE addition prior to infection was not additionally beneficial or detrimental under the investigated experimental conditions. Cell viability was tested in the presence of 5 µM ApoE 2, 3 or 4 confirming that none of the ApoE isoforms affected cell growth (Fig. S3b and S3c). Considering that the phenotype is the same on both SH-SY5Y and GMK, and that GMK is commonly used for HSV1 research in the lab, which gives practical benefits to our study, we decided to work on GMK for the rest of our experiments.

**Fig. 1.**
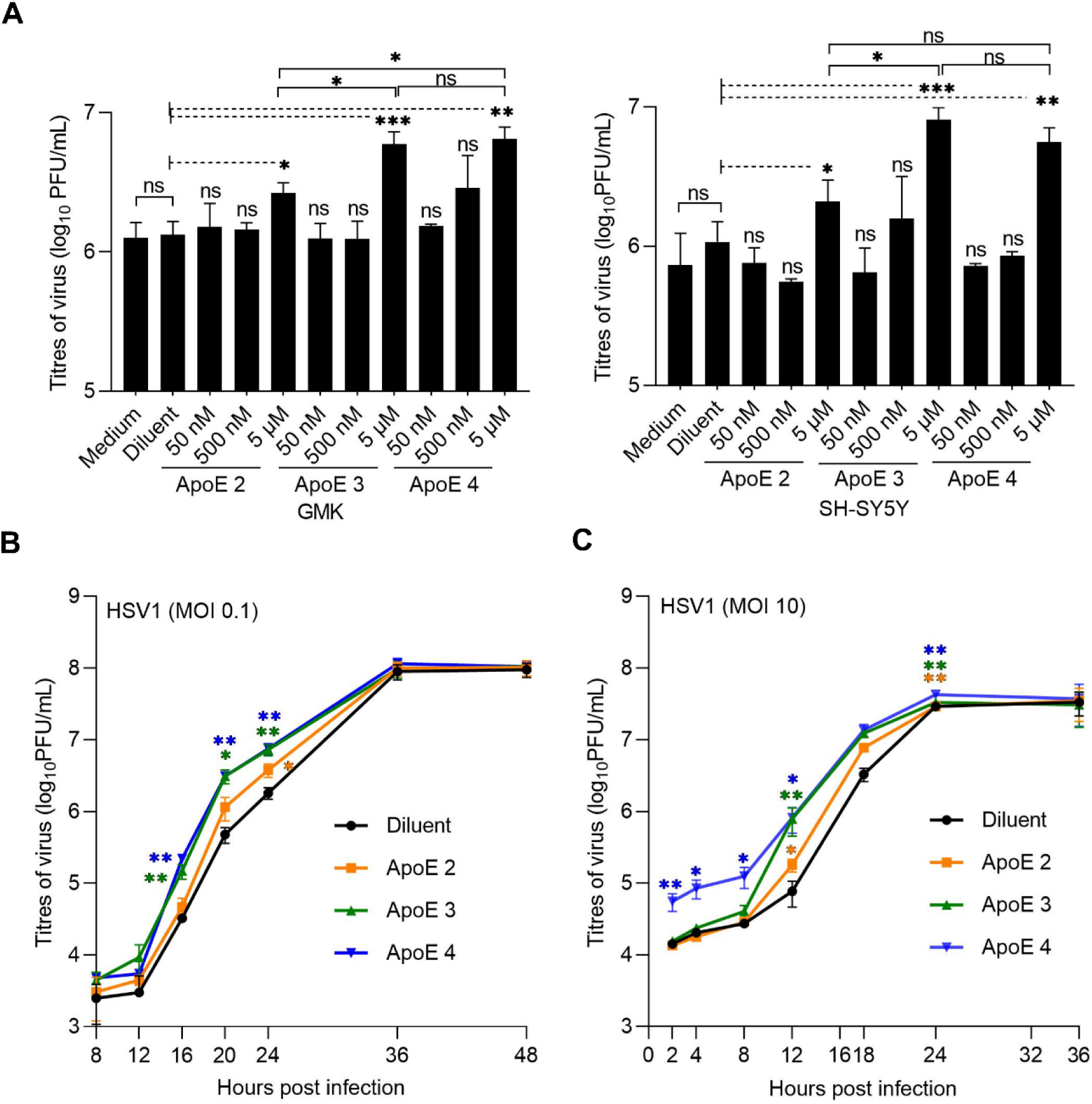
HSV1 growth is accelerated by ApoE. **(A)** HSV1 growth was analysed by plaque assay at various concentrations of different ApoE isoforms on either GMK (left) or SH-SY5Y (right). All the ApoE groups were compared to the diluent group. 5 µM of ApoE 2, 3, and 4 were compared with each other. **(B and C)** HSV1 growth kinetics of multiple (MOI 0.1) **(B)** or single (MOI 10) **(C)** infection cycles were studied on GMK with 5 µM of the different ApoE isoforms. Results represent at least three independent repeats. Error bars: mean ± SD (standard deviation). Student t-test, ns=not significant, *: p≤0.05, **: p≤0.01, and ***: p≤0.001.

Next, we investigated the HSV1 growth kinetics of multiple (MOI 0.1) or single infection (MOI 10) cycles on GMK in the presence of 5 µM ApoE isoforms. During multiple infection cycles (Fig. 1B), HSV1 in all groups had similar titres at shorter time points (< 12 hpi) and reached the same plateau values. However, in comparison to the diluent group, higher virus titres were detected at middle time points (16, 20 and 24 hpi), when ApoE was present (Fig. 1B). Such promotional effects are also isoform dependent, as more viruses were seen with ApoE 3 or 4 than with ApoE 2 at these timepoints (Fig. 1B). Similar as in multiple infection cycles, isoform dependent higher virus titres were also seen at middle timepoints (12 and 18 hpi) for ApoE groups as compared to the diluent group during virus single infection cycle, with more viruses detected in ApoE 3 or 4 group than ApoE 2 (Fig. 1C). An interesting observation was that for the ApoE 4 case, under single infection cycle conditions (high MOI), more HSV1 viruses were detected at early time points, which most likely resulted from the release of extracellular viruses from the cell surface, instead of promoted virus growth, in view of the short time after infection initiation. Considering the similar infection starts and plateaus, these results suggest that one or multiple steps of the HSV1 life cycle are promoted by ApoE, where ApoE 3 or 4 shows stronger effects than ApoE 2. This made us investigate the potential roles of ApoE at different infection steps individually. In our experimental design, purified ApoE was added after 1 h inoculation. Given that most HSV1 can successfully enter cells within the 1h inoculation (Nicola & Straus, 2004), virtually all cells are infected with MOI 10 infection before the addition of ApoE. Thus, no influence of ApoE on binding and entry of the internalized HSV1 is expected for the single infection cycle case (MOI 10, Fig. 1C) and for the first round of infection of the multiple cycle case (MOI 0.1). Thus, to focus on the pro-viral effects of ApoE, we at first directed our attention to the processes after virus entry, including virus replication and egress.

### HSV1 detachment from the cell surface, but not replication, is accelerated in the presence of ApoE

HSV1 replication was analysed by qPCR for genome quantification. At 24 hpi with HSV1 infection of MOI 0.1, supernatant and cell-associated viruses were harvested separately and quantified by plaque assays and qPCR respectively. Titration of the supernatant by plaque assays was included as a control and the results revealed increased viral titres in ApoE added groups (Fig. 2A, left Y axis), in agreement with previous data (Fig. 1). In contrast, cell-associated virus genome copies were similar in all groups (Fig. 2A, right Y axis). These results strongly suggest that ApoE promotes HSV1 release without affecting its replication.

**Fig. 2.**
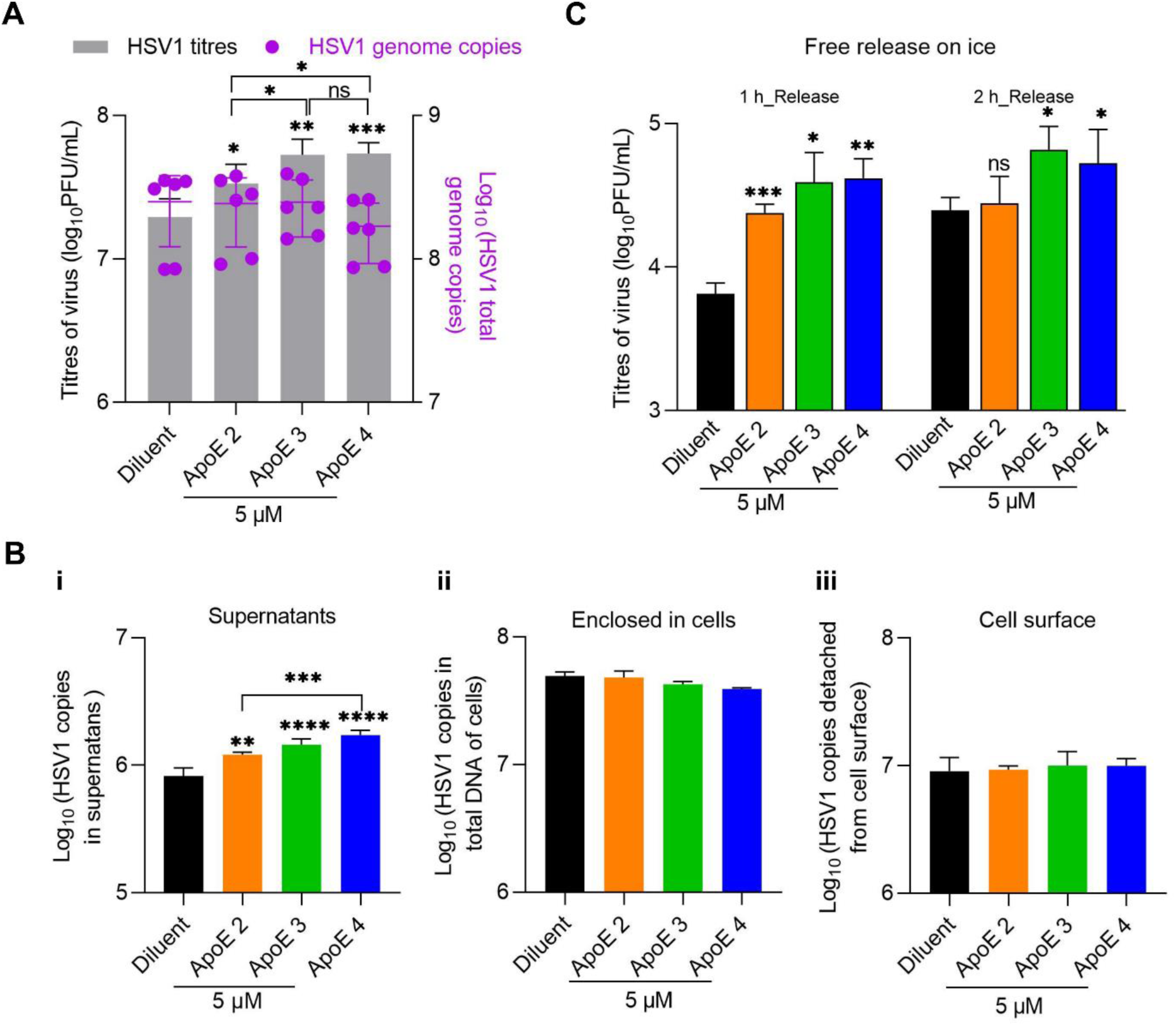
The release of HSV1, but not the replication, is promoted by ApoE. **(A)** GMK cells were infected with HSV1 (MOI 0.1) and ApoE or diluent was added at 1hpi. At 24 hpi, medium and cell samples of different groups were separated and quantified by plaque assay for released viruses (grey bars, left Y axis) and qPCR for cell associated ones respectively (purple dots, right Y axis). **(B)** Infection was done in the same way as in (A). At 20 hpi, HSV1 on the cell surface was released by trypsin after the separation of supernatants (medium) and cell samples as in (A). Supernatants (i), HSV1 released by trypsin (ii, cell surface), or enclosed in cells (iii) were quantified by qPCR. **(C)** Infection was done in the same way as in (A). At 20 hpi, the supernatant was removed, and cells were extensively washed with cold PBS. Virus release was monitored for 1 or 2 h and quantified by plaque assay. For the data analysis in figure (A) and (B), ApoE groups were compared with the diluent individually as well as with each other. The significance is indicated on the top of each column for the virus titres or genomes. No significance was observed for the statistical analysis between any group of genome copies, enclosed cells, or cell surface. Results were from three independent repeats. For the data analysis in figure (C), ApoE groups were compared to the diluent group individually at both time points. Results represent four independent repeats. Student t-test, ns=not significant, *: p≤ 0.05, **: p≤ 0.01, ***: p≤ 0.001 Error bars: mean ± SD.

Next, we quantified the amounts of viruses on the cell surface prior to release. This quantification could be used as an indication for the efficiency of virus assembly and intracellular traffic to the cell membrane. For this purpose, samples were harvested at 20 hpi. After the separation of cells and supernatants, virus particles displayed on the cell surface were released by trypsin digestion and collected. Then the three parts of the sample, i.e, viruses in supernatants, displayed on the cell surface, or enclosed in cells were analysed. Quantification of virus in supernatants by qPCR revealed ApoE isoform-dependent virus promotions (Fig. 2B-i, supernatants), which was also observed by titration of the supernatants via plaque assay (Fig. S4). Quantification of cell-associated viruses did not yield significant differences neither in the cell-enclosed nor trypsin-released (cell-surface) materials, suggesting that virus packaging or trafficking is unlikely modified by ApoE (Fig. 2B-ii and 2B-iii). With more viruses in the supernatant but similar amounts on the surface, our results strongly suggest that HSV1 release from the cell membrane is promoted in the presence of ApoE and that the HSV1 interaction kinetics at the cell surface reach a new equilibrium in the presence of ApoE.

The quantities of HSV1 are similar on cell surfaces of both mock and ApoE treated groups, as shown in Fig. 2B. Taking this as the starting point, we monitored HSV1 release from the cell surface. In this experiment, the supernatant was removed at 20 hpi and the cells were washed with cold PBS. Subsequently, virus release was monitored by harvesting the supernatant at 1 h and 2 h after washing and quantifying its viral titre by plaque assay. As HSV1 trafficking and release are highly dependent on cellular exocytosis (Griffiths et al., 1985), the experiment was performed on ice, in order to inactivate exocytosis, thus excluding the transport of new particles to the cell membrane. Supernatants at 20 hpi were titrated as controls before the evaluation of HSV1 release, which confirmed that there were more viruses in the supernatants of ApoE groups at this time point. For both release time points, more viruses were detected in the supernatants from ApoE groups than that of the diluent group (Fig. 2C), leading to the conclusion that HSV1 detaches faster from the cell surface in the presence of ApoE. As there was no ApoE added in the medium during the release test, we speculate that when HSV1 detaches from the plasma membrane, ApoE is either enriched in the newly produced virus particles or present at the plasma membrane. These suggested scenarios are further investigated below.

### ApoE co-sediments with HSV1 particles

HSV1 has been shown to bind various human lipoprotein particles and artificial proteoliposomes (Huemer et al., 1988). Recent studies have also revealed that ApoE enriches in virus particles produced from ApoE expressing cells and plays important roles for efficient HBV and HCV infections (Bankwitz et al., 2017; Chang et al., 2007; Jiang & Luo, 2009; Qiao & Luo, 2019). Our results made us wonder whether ApoE was also associated with HSV1 after purification, when such particles are produced from an ApoE expressing cell line. To test this, the ApoE expressing cell line Huh-7, permissive to HSV1 infection, was used. Samples from mock or HSV1 infected groups were separated by ultra-centrifugation through layers of sucrose gradients as described in materials and methods. After centrifugation, samples and sucrose gradients were fractionated from top to bottom and processed for detection of ApoE and HSV1 by western blot. The distribution of ApoE, in the mock or HSV1 infected group, shows different patterns across the sucrose gradients (Fig. 3A and 3B). It should be noted that lipoprotein particles associated with ApoE do not penetrate 10% sucrose (Smith et al., 1982); thus fractions 1 to 6 are expected to contain ApoE primarily in particle-free form, the distributions of which were later compared. Without infection, the highest concentration of ApoE was observed in fraction 4, while the concentrations in fractions 1, 2, and 6 were lower and similar to each other (Fig. 3B, ApoE_Mock). In contrast, the distribution of ApoE through the sucrose gradients was modified after infection: the ApoE concentration was the highest in fraction 1 and gradually decreased through the gradients (Fig. 3B, ApoE_HSV1), which is in a similar trend as that of HSV1-gC (Fig. 3B, gC_HSV1), used here as an indicator for which fractions contain virus particles. Titration of the fractionated samples confirmed that detection of HSV1-gC is associated with the presence of infectious HSV1 particles (Fig. 3C). The mismatch between gC (Fig. 3A and B, fraction 3) levels and viral titres (Fig. 3C, fraction 3) indicates the presence of defect or empty viral particles in fractions 1 and 2. The similar distribution patterns of ApoE and HSV1-gC in the infected group strongly indicate that ApoE is hijacked by HSV1 and stays associated with virus particles after release. Further, to estimate the amounts of ApoE that could be associated with HSV1, the distributions of ApoE through the 6 fractions, with or without HSV1 infection were compared, based on the intensities of the corresponding western blot bands from Fig. 3A. In comparison to the distribution of ApoE in the mock group, about 35% of ApoE was shifted after HSV1 infection from fractions 4, 5, and 6 to the top fractions, i.e., fractions 1, 2, and 3. Thus, we propose that the shifted portions (35%) of ApoE is associated with HSV1 after purification.

**Fig. 3.**
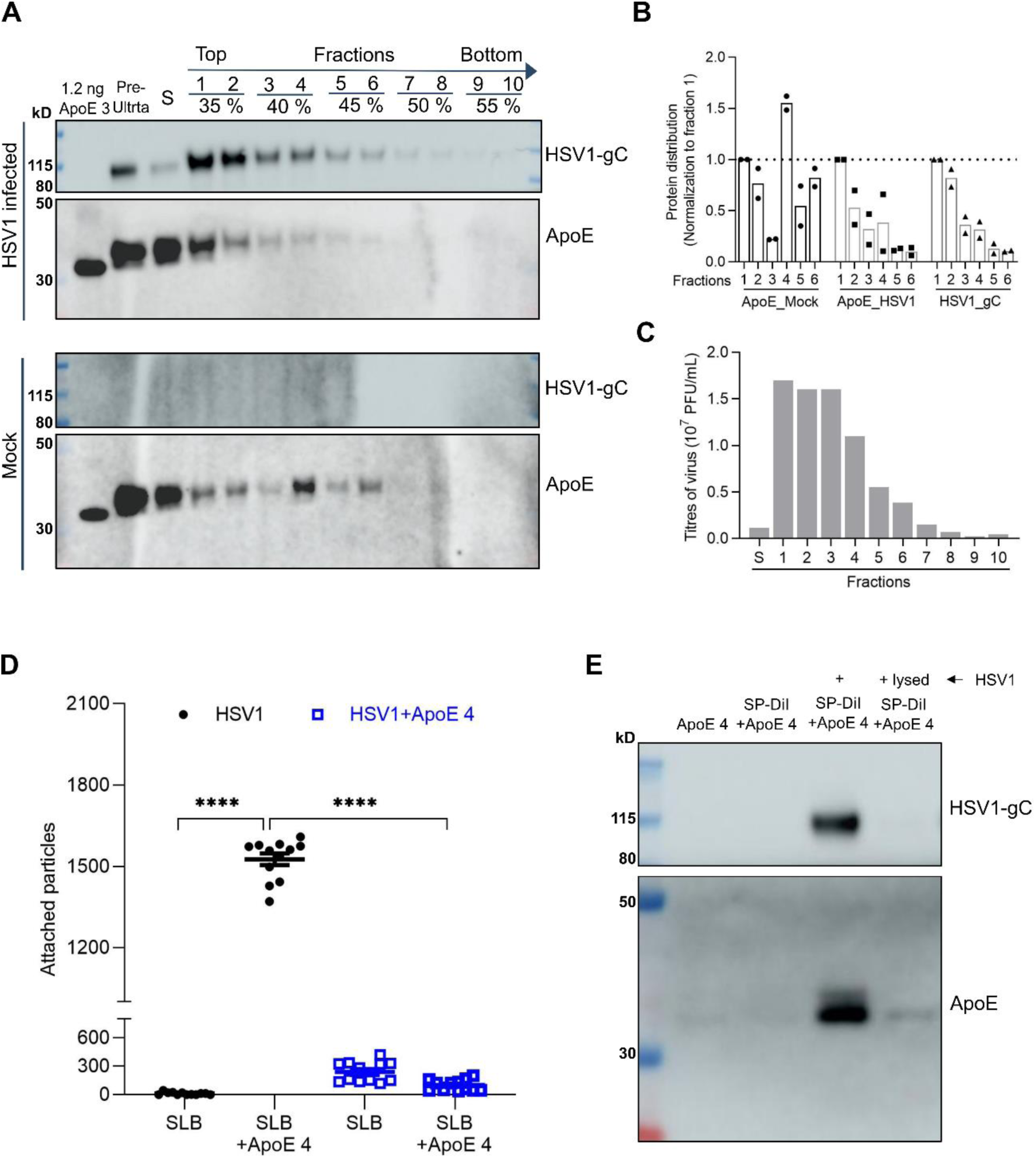
Interactions between ApoE and HSV1 particles. Huh-7 cells were mock or infected with HSV1 (MOI 0.03) for 72 h. Viruses, proteins, and other contents in the supernatants were concentrated and separated through sucrose gradients by ultra-centrifugation. Samples were fractionated from top to bottom after centrifugation. Solution (S) indicates the samples loaded on top of the sucrose gradients after centrifugation. **(A)** Fractions were analysed by western blot and probed for HSV1-gC and ApoE. HSV1-gC was used as a marker to identify the fractions containing virus particles. Purified ApoE 3 (1.2 ng) was included as a control. **(B)** Distributions of ApoE (of mock or HSV1 infected groups) and HSV1-gC from fractions 1 to 6 were displayed based on the intensities of the corresponding blots from (A). The intensities of different fractions were normalized to the value of fraction 1. Results were generated from two independent repeats. Both the individual values (round, square, or triangle dots) and their means (bars) are displayed for each fraction. **(C)** HSV1 titres of the corresponding fractions (HSV1 infected groups) were quantified by plaque assays. Results are shown from one of the repeats. **(D)** Numbers of HSV1 and ApoE 4 coated HSV1 particles on a POPC:DGS-NTA membrane surface with or without His-ApoE 4 immobilized. 2×10^8^ PFU HSV1, 4 µM ApoE, and 40 µM fluorescent lipophilic dye (SP-DiI) were incubated for 1h in PBS and separated as described in the material and method. Fluorescently labelled viral particles were analysed for their attachment to immobilized His-tagged ApoE 4. Each point in the plot represents particle count from a single image of 6 randomly selected fields. Data were produced from 2 independently prepared surfaces. Statistics done by one sample t-test. Error bars indicate mean and standard error of mean (SEM). ****: P≤ 0.0001. **(E)** Samples after incubation with the lipophilic dye and ApoE and purification were lysed and processed for western blot. HSV1-gC and ApoE were probed.

Our results are in line with the previously described interactions between HSV1 particles and lipoprotein particles (Huemer et al., 1988). However, if there are interactions between virus particles and purified ApoE protein, and whether ApoE can associate with the virion after particle formation remains unanswered. To test this, direct interactions were analysed by total internal fluorescence microscopy (TIRFM) visualization of fluorescently labelled HSV1 particles binding to surface-immobilized ApoE. As shown in Fig. 3D, HSV1 binding was dramatically increased when ApoE 4 was present (Fig. 3D, black dots), demonstrating direct interactions between HSV1 and ApoE 4. The apparent affinity of this interaction was found to be isoform-dependent, with ∼3 times higher apparent affinity of ApoE 4 to the virus particle as compared to ApoE 2 and ApoE 3 mostly driven by an increase in the association rate constant of the particle to the protein (Fig. S5). To further strengthen our results, we tested whether ApoE 4 added exogenously can bind stably to HSV1 particles and thus be co-purified together with HSV1 as complexes after pre-mixing the two. In this case, a purification protocol based on size-exclusion was developed to efficiently remove free protein while recovering intact particles. Western blot analysis showed that ApoE 4 was recovered after purification only in presence of intact HSV1 particles, confirming that the protein interacts with the virus particles when added exogenously (Fig. 3E). Lysed HSV1 was included as a control, as the separation is based on the sizes of complexes. In this case, neither HSV1 nor ApoE were collected by during purification procedure (Fig. 3E). Finally, it was shown that the binding of ApoE 4-coated HSV1 particles to surface-immobilized ApoE 4 decreased below control levels, further confirming an interaction between HSV1 particles and ApoE 4 (Fig. 3D, blue squares). From these results, we conclude that purified ApoE 4 can interact with HSV1 particles, and that the complexes formed by ApoE and HSV1 remain stably associated after separation of free ApoE.

### ApoE decorated HSV1 demonstrates faster attachment and higher efficiencies of both attachment and entry to cells

The observation that ApoE associates with HSV1 particles raises the question of whether ApoE associated with HSV1 modifies the interactions between HSV1 and the cell surface thereby influencing the attachment and entry process. This was tested by comparing the binding and entry of HSV1 and ApoE 4-coated HSV1 separately. To get the same input for both types of HSV1 during the binding experiment, freshly prepared viruses were quantified by qPCR and the same quantities (50,000 copies) of each were added to cells for attachment on ice. Our results show that there were more copies of ApoE 4-coated HSV1 attached to the cell surface than for the ApoE-free HSV1 group at most time points (Fig. 4A), which indicates faster attachment kinetics and higher attachment efficiencies. At 60 min after attachment on ice, the quantities of ApoE 4-coated HSV1 are ∼2 times higher than that of ApoE-free HSV1. To compare the entry behaviour, while taking into account differences in attachment efficiency, the amounts of entered viruses (the plaque numbers) were normalized according to the amount of particles bound to the cell surface. These binding quantities were determined by qPCR quantification of the number of attached particles on ice, parallel to the entry experiment (see details in the methods). As displayed in Fig. 4B after normalization, ApoE 4-coated HSV1 demonstrated slightly higher entry quantities at all time points, (Fig. 4B), although the difference was only statistically significant 60 min after entry initiation. Taken together, our results indicate that harbouring ApoE can also benefit to HSV1 by facilitating virus attachment and entry into cells.

**Fig. 4.**
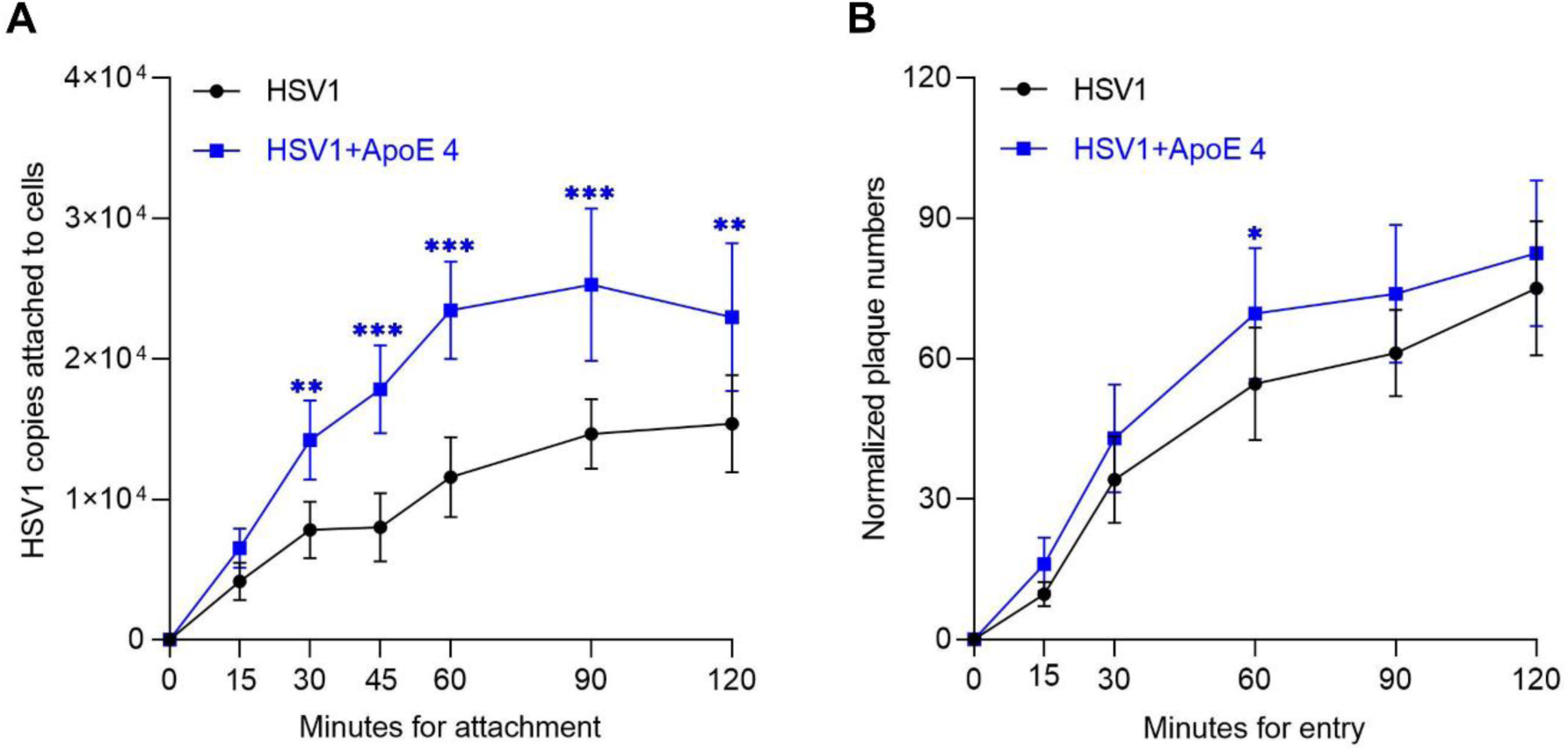
Comparison of the binding and entry of HSV1 and ApoE4 coated HSV1. **(A)** The same amounts (50,000 copies of each according to the qPCR quantification) of the freshly prepared HSV1 and ApoE 4-coated HSV1 (as in Fig. 3E) were added to GMK cells for attachment on ice. At indicated times, the attached viruses were collected and quantified by qPCR. The data represent results from three independent repeats. **(B)** The entry kinetics of HSV1 or ApoE 4-coated HSV1 was studied after 1h synchronization on ice. Before entry initiation, the amounts of attached virus after 1h synchronization on ice were quantified by qPCR in a parallel group. The plaque numbers are presented after normalization to the amounts of attached viral copies. Results are the summary of three independent repeats. Student t-test, significance *: p≤ 0.05, **: p≤ 0.01, ***: p≤ 0.001 Error bars: mean ± SD.

### The interactions between HSV1 and the cell surface are modified by virus-bound ApoE

The results of the facilitated attachment, entry, and release of HSV1 suggest modified interactions between HSV1 and the cell plasma membranes by ApoE. To further characterize these interactions, we use a well-controlled biophysical experimental system, which allows us to focus on the effect of ApoE on the virus-membrane interaction, while excluding the contribution of other cellular factors. In this experiment, we characterized the interaction kinetics of HSV1 particles decorated with ApoE or ApoE-free HSV1 particles, to membranes from GMK cells, using TIRFM. For this purpose, native supported lipid bilayers (nSLBs) (Pace et al., 2015; Peerboom et al., 2018), i.e. two-dimensional planar lipid membranes supported onto a sensor substrate and containing plasma membrane material, were used as cell-surface mimetics in biophysical studies of virus membrane interactions. TIRFM allows for the visualization of surface-bound viruses while discriminating them from the ones in solution (Fig. 5A). This makes it possible to analyse virus-membrane interaction kinetics on a single particle level. The association of viruses to the surface can be plotted over time (Fig. 5B) and the arrival rate, directly proportional to the association rate *k*_on_, can be quantified, provided the experiment is carried out under reaction limited conditions. Additionally, information on the dissociation behaviour of the virus from the membrane can be obtained by analysis of the particle residence time; fitting of the dissociation curves with a double exponential decay function with an offset (Fig. 5C) allows for a quantification of the apparent dissociation rate constants (*k*_off_) (Fig. 5D) (Peerboom et al., 2018) as well as of an irreversible fraction determined from the offset value (see the materials and methods for details). These experiments revealed that ApoE 4-coated HSV1 associated significantly faster to the membrane, showing an average 4.4 ± 2.6 times faster arrival rates than noncoated HSV1 (Fig 5D-i). The virus also exhibited a heterogeneous detachment behaviour; virus dissociation could broadly be separated into two dissociating populations characterized by fast (*k*_off_^1^) or slow (*k*_off_^2^) dissociation rates and a population of irreversibly bound (non-dissociating) particles. On average, 85% (ApoE 4-free) and 77% (ApoE 4-coated) of the particles were bound irreversibly (Table S1) while the fit revealed a roughly equal fraction of the remainder dissociating particles belong to *k*_off_ ^1^ resp. *k*_off_ ^2^, in all cases (Table S1). The values of *k*_off_ ^1^ were on average 0.023s^-1^ for HSV1 and 0.027s^-1^ for ApoE 4-decorated HSV1: they were 0.0010s^-1^ and 0.0011s^-1^ respectively for *k*_off_^2^ (Table S1). The values for the slow dissociating fraction are comparable with what observed in similar experimental setup for the interaction of herpes simplex virus to GMK cells (Peerboom et al., 2018). From these values we can estimate the unbinding energy for the two subpopulations assuming for simplicity a single energy barrier with logarithmic kinetics (Bally et al., 2011). Under these assumptions, the binding energy (*E*_B_*^n^*) can be calculated from *k*_off_*^n^*. as:

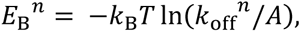

**Fig. 5.**
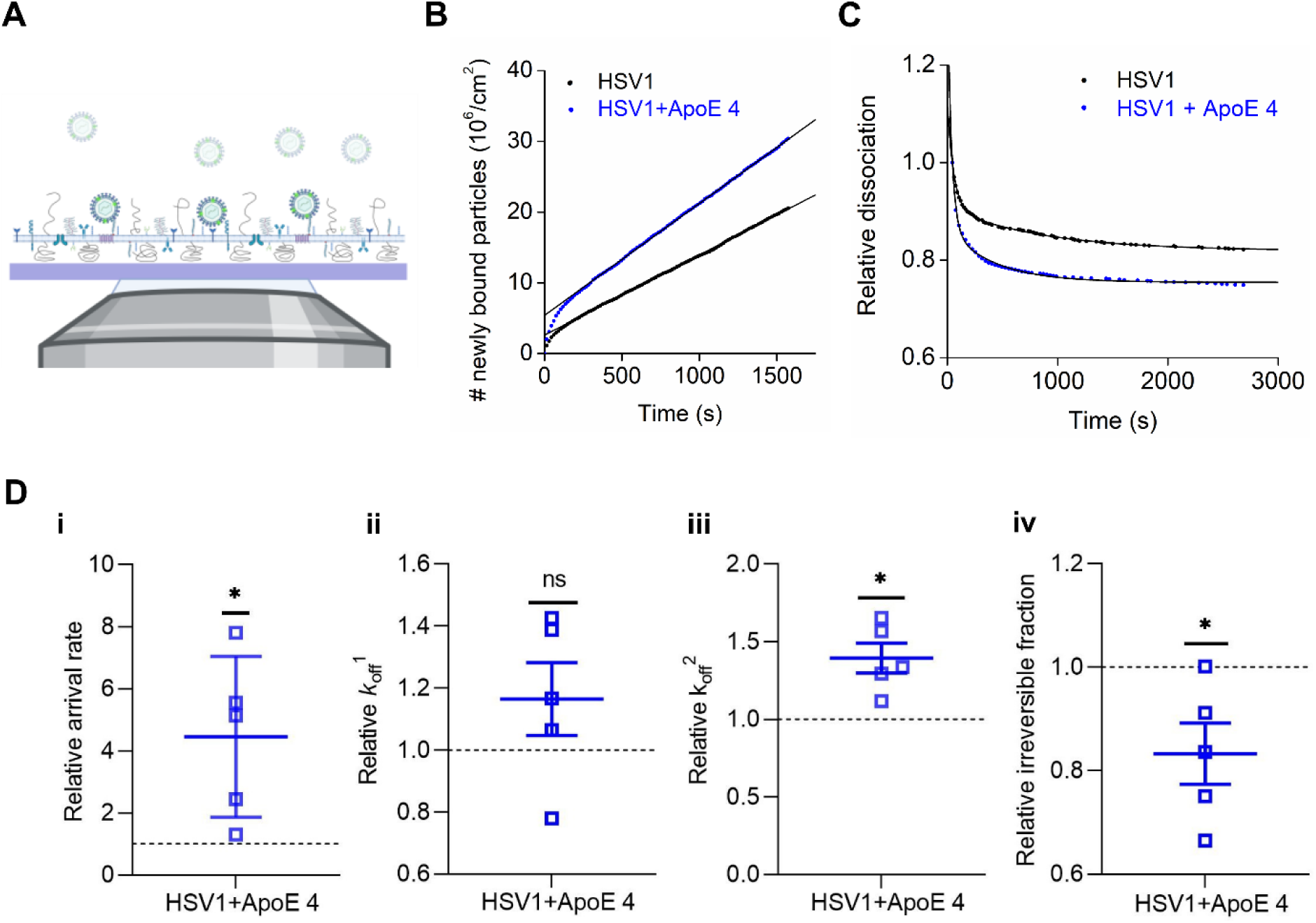
ApoE 4-coated HSV1 particles display faster attachment and detachment from the native supported bilayers. **(A)** Schematic representation (not to scale) of TIRFM-based assays to probe the interaction of fluorescently labelled particles to nSLBs. Planar nSLBs are formed on a glass substrate, before being incubated with fluorescent viruses. The interaction between the fluorescent viruses and the nSLBs from GMK membranes (lacking ApoE) can then be imaged using TIRFM, selectively illuminating only viruses interacting with the surface, while ignoring free viruses in the bulk solution. Representative curves of **(B)** the association and **(C)** the dissociation kinetics of HSV1 (black) and ApoE 4-coated HSV1 (blue) to nSLBs with the corresponding linear and double exponential decay fits, respectively. **(D)** Mean and SEM of the relative association (D-i, arrival rate) from the linear fit of curves as shown in (B) and relative dissociation kinetics (D-ii, *k*_off_^1^; D-iii, *k*_off_^2^; and D-iv, irreversible fraction) from the double exponential fit with an offset, of the data as shown in (D) (see materials and methods for details). Blue squares represent the binding data for ApoE 4-coated HSV1 particles to nSLBs, which are normalized to HSV1 data (black dotted line). Each data point is a sum of particles at 3 different positions per well from at least 2 independent experiments. Statistical significance is calculated using one sample t-test with mean 1 (*: p ≤ 0.05). Schematic in panel (A) was made with BioRender.com.

Where *k*_B_ is the Boltzmann constant, T is the absolute temperature and A is a constant indicating the attempt rate and usually in the order of 10^13^ s^-1^. The value of *E*_B_^1^ is in the order of 83 kJ/mol while *E*_B_^2^ is 90 kJ/mol. Thus, the energy reduction due to the presence of ApoE, which is independent of the value of A, is 0.37 kJ/mol for the fast component and 0.88 kJ/mol for the slow one.

Although no significant difference was observed for the fast dissociation rate constant (Fig. 5D-ii, *k*_off_^1^), ApoE 4-decorated HSV1 particles were found to exhibit 39% higher dissociation rates for the slower dissociation rate constant compared to HSV1 (Fig. 5D-iii, *k*_off_^2^). The ApoE 4-decorated particles also show an 8.3% reduction of irreversibly bound particles compared to HSV1 (Fig. 5D-iv, irreversible). Combined, these results indicate that ApoE-decoration of HSV1 enhances the dynamics of virus-membrane interaction by increasing both the association and dissociation of viral particles from the membranes.

### ApoE 4-coated HSV1 exhibit increased association and dissociation to heparan sulfate

In view of the reported interaction between ApoE and HS (Saito et al., 2003), we hypothesized that the modified interactions between HSV1 and cell plasma membrane resulted from the interactions between HSV1 and HS on the cell surface. HS is abundantly expressed on the surface of GMK cells (Feyzi et al., 1997) and also found in nSLBs derived from such cells (Peerboom et al., 2018). The hypothesis was tested by looking at the binding of HSV1 and ApoE 4-coated HSV1 particles to HS. Using TIRFM measurements on surface-bound HS (Fig. 6A-C), the ApoE 4-coated particles showed on average 2 times higher association rate to HS as compared to non-coated particles (Fig. 6D-i). Although both ApoE 4-coated HSV1 and HSV1 particles detached with similar fast (*k*_off_^1^) and slow (*k*_off_^2^) dissociation rates from HS (Fig. 6D-ii and 6D-iii), a significant reduction (28%) of irreversibly bound particles was found for ApoE 4-coated HSV1 as compared to ApoE-free HSV1 (Fig. 6D-iv).

**Fig. 6.**
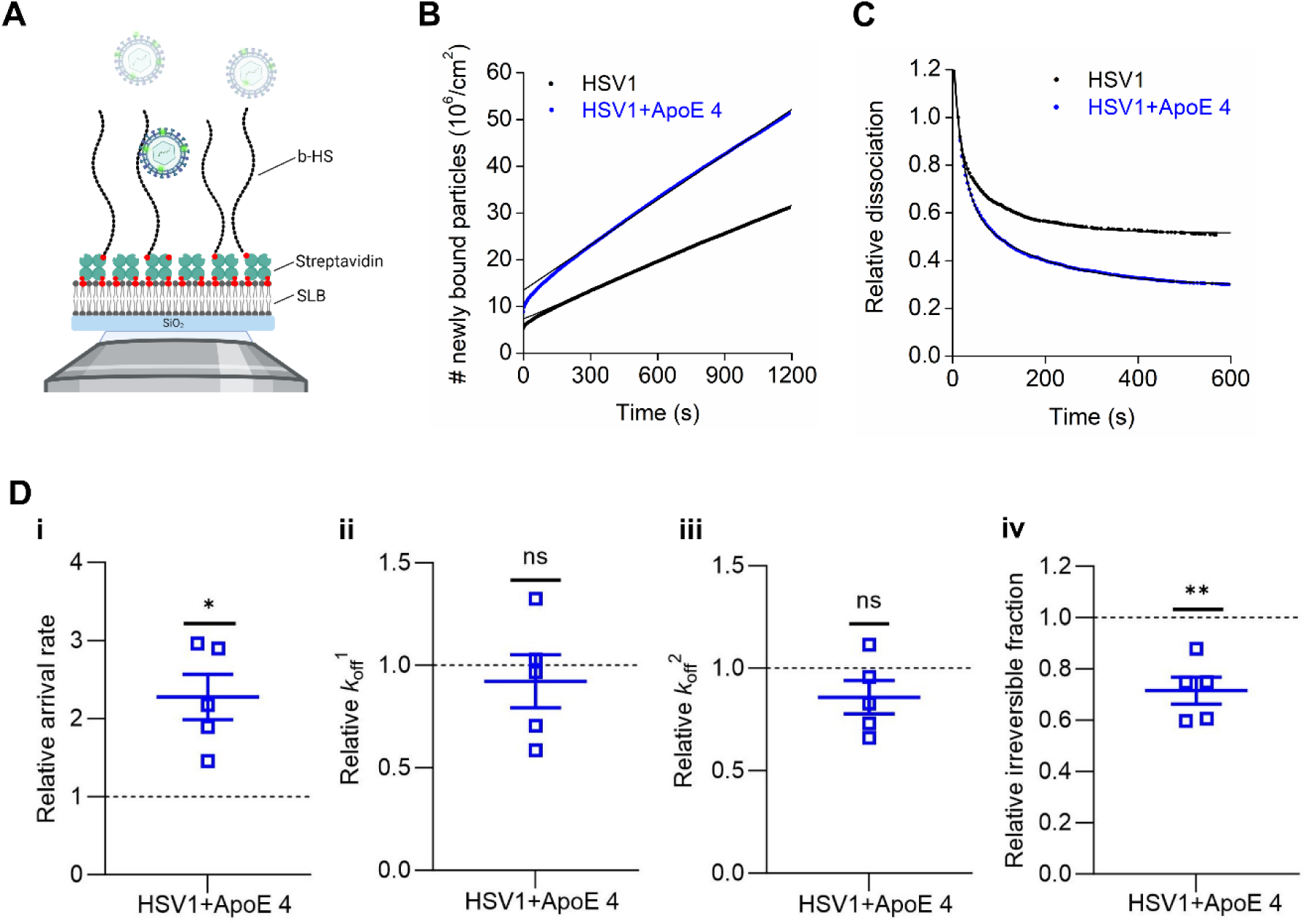
ApoE 4-coated HSV1 attaches faster to heparan sulfate. **(A)** Schematic representation (not to scale) of TIRFM-based assays to probe the interaction of fluorescently labelled particles to surface-bound HS on SLBs using streptavidin layer as a sandwich between SLBs and biotinylated HS (b-HS). Representative curves for **(B)** the association and **(C)** the dissociation kinetics of fluorescently labelled HSV1 (black) and ApoE 4-coated HSV1 (blue) particles to HS films with the corresponding linear and double exponential fits, respectively. **(D)** Mean and SEM of the relative association kinetics (D-i, arrival rate) from the linear fit of the curves as shown in (B) and relative dissociation kinetics (D-ii, *k*_off_^1^; D-iii, *k*_off_^2^; and D-iv, irreversible fraction) from the double exponential fit with an offset of the data as shown in (C) (see materials and methods for details). Blue squares represent the binding data for ApoE 4-coated HSV1 particles to HS, which are normalized to HSV1 data (black dotted line). Each data point is a sum of particles at 3 different positions per well from 3 independent experiments. Statistical significance is calculated using one sample t-test with mean 1 (*: p ≤ 0.05 and **: p ≤ 0.01). Schematic in panel (A) was made with BioRender.com.

Altogether, these results indicate that ApoE 4 promotes both association and dissociation of HSV1 on HS films. In conjunction with the nSLBs results (Fig. 5), this implies that the presence of HS in nSLBs might be a dominant factor regulating the interaction between HSV1 and HS through ApoE.

### HSV1 particles have similar dissociation rate constants (***k*_off_**) on membranes with or without ApoE association

We also hypothesized that HSV1 would detach faster from membranes harbouring ApoE. The residence of ApoE in the plasma membrane, together with caveolin, has been reported in adipocytes (Yue & Mazzone, 2011). Thus, we first tested whether ApoE was also membrane-associated in cells of relevance to our experiment. For this purpose, an ApoE 4 expression inducible cell line was established. The induction of ApoE 4 expression was verified by western blot (Fig. S6a); and the presence of ApoE 4 was also confirmed in the plasma membrane materials isolated after sucrose purification (Fig. S6b). To compare the influence of membrane-bound ApoE on virus binding kinetics, we produced nSLBs, from both ApoE 4-induced and non-induced cells. Kinetic analysis (Fig. 7A and B) with TIRFM revealed that the presence of ApoE in the membrane does neither influence the association (Fig. 7C-i) nor the dissociation behaviour (Fig. 7C-ii, iii, and iv) of the virus from the membrane with or without the presence of ApoE. These results indicate that membrane associated ApoE is not a significant factor in promoting HSV1 attachment or release.

**Fig. 7.**
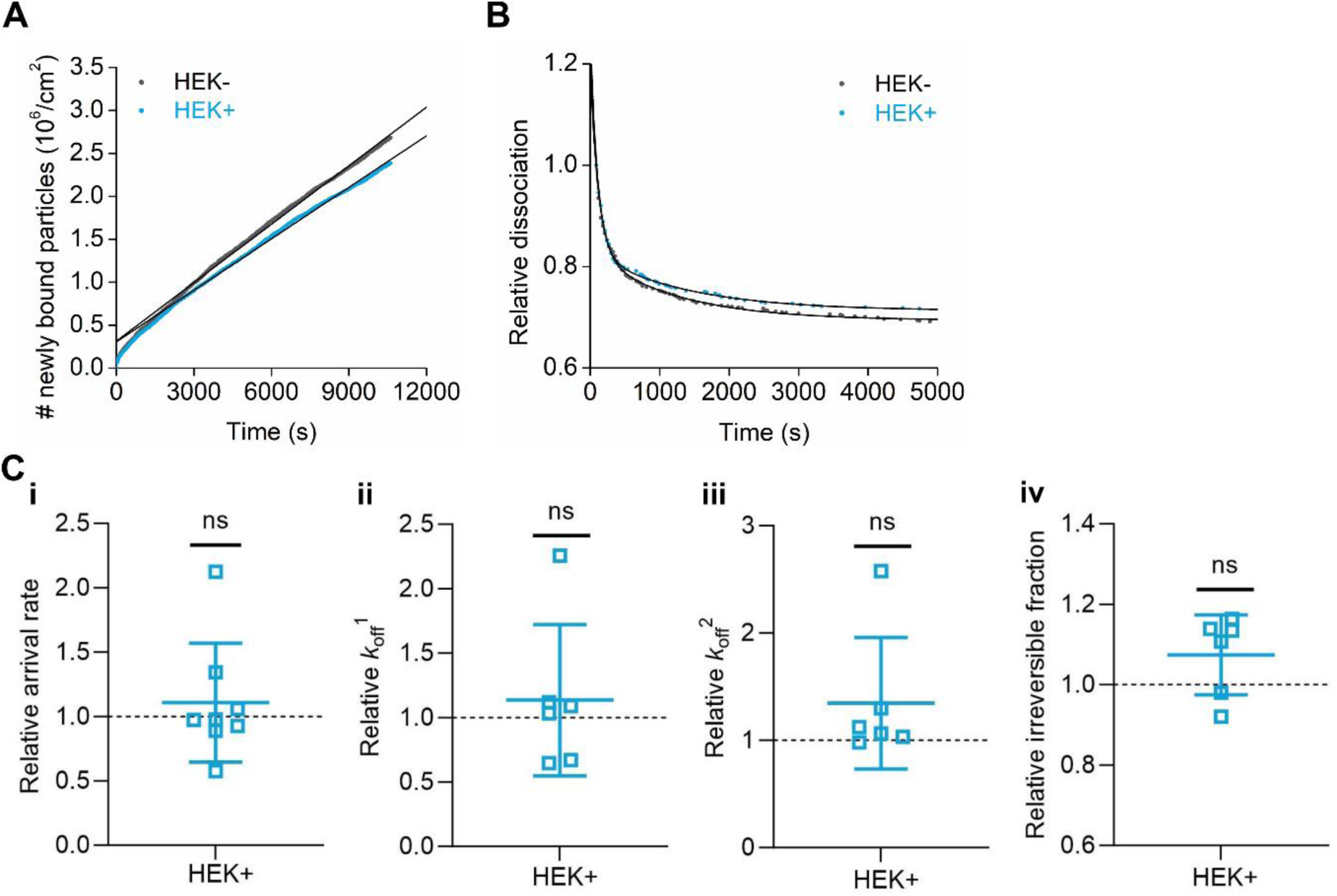
HSV1 detaches with similar *k*_off_ on native supported lipid bilayers in the presence or absence of ApoE. Representative curves of **(A)** association and **(B)** dissociation kinetics of fluorescently-labelled HSV1 from nSLBs generated from the membrane materials of HEK cells without (HEK-, black) or with (HEK+, cyan) the induction of ApoE 4, with their corresponding linear and double exponential fits, respectively. **(C)** Mean and SEM of the association kinetics (C-i, relative arrival rate) from the linear fit of the data as shown in (A) and dissociation kinetics (C-ii, iii, and iv, relative *k*_off_^1^, *k*_off_^2^ and irreversible fractions, respectively) from the double exponential fits with an offset of the data as shown in (B) (see materials and methods for details). Cyan squares represent the binding data of HSV1 particles to nSLBs with the induction of ApoE 4 (HEK+), which are normalized to the binding data of HSV1 particle to nSLBs without the induction of ApoE 4 (HEK-, black dotted line). Each data point is a sum of particles at 3 different positions per well from three independent experiments. Statistical significance is calculated using one sample t-test with mean 1 (ns = not significant).

## Discussion

A number of epidemiological studies have reported that HSV1 infection and ApoE associate with a higher risk of AD (Itzhaki et al., 2001; Itzhaki et al., 1997; Lin et al., 1996; Linard et al., 2020; K Lopatko Lindman et al., 2019; Lövheim et al., 2019; Steel & Eslick, 2015; Strandberg et al., 2005). Thus, we studied potential HSV1-ApoE interactions during virus infection cycles on a molecular level. In this study, we monitored HSV1 growth on cultured cells in the presence of different ApoE isoforms and analysed the effects of ApoE on different stages of the HSV1 infection cycle. HSV1 infection was evaluated after adding exogenously expressed and purified ApoE to GMK and SH-SY5Y cells with negligible ApoE expression (Fig. S1a). Such an experimental setup reflects HSV1 infection in neurons, as neurons are suggested to take up ApoE secreted by astrocytes or macrophages in the brain (Boyles et al., 1989; Herz & Bock, 2002). Overall, we found that HSV1 propagation is promoted when it is grown in the presence of ApoE. Our detailed investigations further demonstrate that HSV1 interacts with ApoE, and that this association facilitates both attachment to and release from the plasma membrane, thereby explaining the observed proviral effect.

The promotion of HSV1 by ApoE is concentration-dependent and was primarily observe in our system at concentrations in the micromolar range (Fig. 1A and B). Such a dependence is likely related to the uptake efficiency of ApoE by cells in our artificial *in vitro* system, as it was shown that ApoE addition at 2.5 µM or 5 µM is needed to guarantee an effective uptake of the protein by most cells, which is highly correlated to the promoted HSV1 growth at these two concentrations (Fig. S2). As the primary goal of this project was to investigate the potential influence of ApoE on HSV1 infection, we decided to continue with 5 µM ApoE that ensures a widespread distribution of the protein in most cells in our experimental system. Importantly, cell metabolism was not altered when this concentration of ApoE was added (Fig. S3c and d), which implies that the promotion of HSV1 is not an artifact or an indirect effect resulting from a change in cell behaviour. It should also be noted that ApoE concentrations in the medium of Huh-7 cells in culture (Fig. S7), the cell line commonly used for studies on hepatitis viruses and ApoE, are in the same range as the concentrations of exogenous ApoE added here, further supporting the relevance our choice. A broad range of physiological concentrations of ApoE, have been reported in the literature, depending on the body fluid type as well as on the applied measurement technique. Reported concentration values typically range from 0.1 to 1.75 µM (Krastins et al., 2013; Hansson, et al., 2014; Nielsen, et al., 2014; Rasmussen et al., 2016; Rezeli et al., 2015; Simon et al., 2012; Wang et al., 2012; Wildsmith et al., 2009), although it cannot be excluded that ApoE concentrations are higher locally. Moreover, local ApoE levels can change in response to stimuli e.g in the case of injury (Boyles et al., 1990; Müller et al., 1985), which has been suggested as a strong epigenetic risk factor for the development of AD (Fleminger et al., 2003; Plassman et al., 2000). After brain injury, accumulation of ApoE in the regeneration sites can last over a month, and the peak of ApoE increase can reach over 250-fold (Boyles et al., 1990). It is thus compelling to speculate that increased ApoE concentrations after traumatic brain injury would be beneficial to HSV1 growth, which may accelerate disease development in HSV1 infected individual.

Incorporation of ApoE into virus particles has been reported for HBV and HCV (Gastaminza et al., 2010; Qiao & Luo, 2019), with consequences on the virus attachment and detachment behaviour. Regarding the attachment of HBV or HCV, ApoE has been suggested to work as a bridge ligand between virus and cell surface receptors, including HS (Jiang et al., 2013; Li & Luo, 2021; Xu et al., 2015), the virus itself lacking HS-binding capacity. In addition of modifying virus interactions at the cell surface by affecting attachment to HS, ApoE has also been proposed to be involved in the late steps of HCV infection (Bankwitz et al., 2017; Benga et al., 2010; Cun et al., 2010; Lee et al., 2014), although whether HCV release is affected by ApoE is unknown. Our study reports a direct biomolecular interaction between ApoE and HSV1, a conclusion supported by two sets of data generated using distinct methods: co-sedimentation of ApoE and HSV1 after virus harvesting from ApoE-producing Huh-7 cells and ultracentrifugation (Fig. 3A); and purification of ApoE-associated HSV1 after incubation of the virus with exogenous ApoE (Fig. 3D and E). These results are complementary to a previous study reporting interactions between HSV1 and purified serum lipoparticles or artificial proteoliposomes harbouring apolipoproteins, including ApoE (Huemer et al., 1988). ApoE distributes across the cytoplasm and the cell plasma membrane both in ApoE-producing Huh-7 (data not shown) and when ApoE is added exogenously to GMK cells (Fig. S9a). It can therefore be inferred that HSV1 may accumulate ApoE after its replication and packing in the nucleus, i.e., during viral particle envelopment, cytoplasmic trafficking, budding on cell plasma. Association can also occur after virus release and such an association with exogenous ApoE appears to be stable over several hours as concluded from the lack of binding of ApoE-coated particles to surface-immobilized ApoE (Fig. 3D, SLB+ApoE 4), even several hours after purification of the complexes. Interestingly, this association was found to be isoform-dependent, with ∼3-fold increase of attachment rate for ApoE 4 to the virus particle as compared to ApoE 2 and ApoE 3 (Figure S5). An interesting, yet still unanswered question, concerns the viral biomolecules mediating the interactions between ApoE and HSV1 particles. Such interactions are likely mediated by viral component(s) on the surface of virus particles, either viral glycoproteins or the lipid membrane of the virus. From our attempts at revealing the molecular binding partner on HSV1, we have excluded potential interactions between ApoE and some HSV1 glycoproteins, including glycoprotein B, C, D, and E, by immunoprecipitation test (Fig. S8). These results indicate that other glycoproteins should be investigated as potential ApoE interaction partners in future studies, or that the protein interacts directly with the lipid envelope.

A significant promotion of virus particle production in the presence of ApoE was observed both in single (Figure 1D) and multiple cycle (Figure 1C) experiments. Noteworthy, is that the growth benefits are isoform-independent, with higher efficiency for ApoE 3 and 4, somewhat in agreement with the ApoE association to HSV1 particles and in contrast to what observed for HCV (Cun et al., 2010). Initial infection experiments were carried out by adding ApoE after 1 hour virus inoculation, followed by rinsing, an experimental setup which does not consider a putative effect of ApoE on virus attachment and entry during the first round of infection, given that HSV-1 entry into GMK cells is expected to occur within minutes (Nicola & Straus, 2004). This indicates that the pro-viral effect is likely ascribed to later steps in the virus replication cycle. Careful investigation of the different steps leading to the production of progeny viruses reveals that ApoE significantly promotes virus release. Indeed, both experiments on live cells following the release of HSV1 into the supernatant in absence of exocytosis (Fig 2C), as well as biophysical experiments detailing the interaction kinetics of individual virus particles with isolated plasma membranes (Fig 5D), reveal that virus detachment is greatly facilitated by the presence of ApoE on the virus particle. The release of HSV describes the detachment of viral particles from the cell surface, which is the final step of the virus infection cycle during its cell-free transmission. In general, this process is much less studied in comparison to other steps. Several studies on this topic have pointed to the effects of viral proteins (Altgärde et al., 2015; Park et al., 2015; Trybala et al., 2021) and/or host factors (Hadigal et al., 2015b; Hopkins et al., 2018a) on virus release. Most of these studies stress that efficient HSV release relies on fine-tuning and overcoming the interactions between viral proteins and GAGs on the cell surface. This led us to speculate that ApoE may influence HSV1 release, by acting as a modulator of virus-GAG interactions, a hypothesis well in line with our observation that ApoE-carrying HSV1 also dissociates faster from isolated surface-immobilized HS molecules (Fig. 6D). While the molecular mechanism behind this increased dissociation remains to be elucidated in detail, we speculate that addition of ApoE to the virus particle influences the multivalent interaction by interfering, possibly through steric hindrance, with the creation of high affinity bonds with the virus glycoproteins (Herold et al., 1991, 1994).

Alpha herpesvirus particles are transported in membrane vesicles (reviewed in (Ahmad & Wilson, 2020)) to specific sites on the cell surface (Hogue et al., 2014; Mingo et al., 2012) via cellular exocytosis pathways. The actual delivery is completed via membrane fusion between the virus-containing vesicles and the cell plasma membrane. It is unknown whether the final release happens at the same sites where virus particles become exocellular or at different sites which requires lateral movement/transport of extracellularly exposed virus particles from the membrane fusion sites. In either case, the potential interventions from ApoE or other host factors should occur at the actual virus release sites. With this assumption, it can be speculated that hijacking ApoE by the virus would be a smarter solution to localize proper amounts of ApoE at the release site than having ApoE homogeneously distributed in the plasma membrane. This may explain why we have observed improved virus detachment with ApoE coated HSV1 (Fig. 5), but not for non-coated HSV1 on nSLBs harbouring ApoE (Fig. 7). It is worth mentioning that the nSBLs were generated from non-infected cell materials (HEK or HEK-ApoE). Thus, our tests cannot exclude the possibility that virus infection modifies and re-arranges membrane association of ApoE in a way that facilitates final release.

The realization that ApoE binds stably with HSV1, even when added exogenously, prompted us to further investigate whether this association can contribute, positively or negatively, to the observed proviral effect. Such an influence becomes more important in the multiple infection cycle experiment (Fig 1B), where ApoE-coated progeny virions are involved in subsequent cellular infection cycles. Virus attachment and entry experiments reveal that the presence of ApoE on HSV1 promotes virus attachment to and entry into cells (Fig. 4). Promoted attachment is further confirmed by biophysical experiments revealing an increased association rate constant (*k*_on_) for ApoE-coated particles as compared to the naked ones. This effect is also likely to be related again to the initial interaction of the virus with HS on the cell surface, which is a known receptor for both ApoE (Saito et al., 2003) and HSV1 (Shieh et al., 1992). Indeed, harbouring ApoE can provide extra ligands to HS for HSV1 attachment to cells on the premise that the viral ligands to HS are not overlapping with the ones to ApoE. This appears to be the case for HSV1 since both gB and gC of HSV1 have been proposed to bind HS (Herold et al., 1991, 1994) but have not been found to mediate HSV1 association of ApoE (Fig. S8). As a result, we propose that ApoE-coated HSV1 has a higher binding avidity to HS due to an increased number of binding sites and that this gives the virus advantages of the faster and more efficient attachment to cells (Fig 4). This idea is strongly supported by the kinetic studies of HSV1 binding to purified HS, where we observed faster association of ApoE 4-coated HSV1 to the immobilized HS (Fig 6D). Taken together, with the faster detachment discussed above, we speculate that the presence of ApoE significantly impacts the molecular characteristics of the interaction between HSV1 and GAGs by increasing the number of binding sites, while reducing the average affinity of the single bonds formed.

When considering the subsequent infection rounds in the multiple infection cycle experiment, it is important to realize that the presence of HS-bound ApoE on the cell surface, may under specific conditions, also lead to virus binding inhibition, and prevent cell-free virus transmission. Such an effect has been reported for example by Dobson and colleagues who show that dimers of a peptide derived from ApoE’s HS-binding domain clearly inhibit virus attachment and subsequent infection of HSV1 and other viruses at high concentrations (>15 µM) (Dobson et al., 2006). Along these lines, we also observed that addition of ApoE 4 h prior (Fig. S9a) to HSV1 infection followed by removal of ApoE from the medium prior infection, leads to significant inhibition of binding (Fig. S9b), but not entry kinetics or efficiency (Fig. S9c and d). We speculate that high concentrations of ApoE or adding ApoE before HSV1 infection can lead to a reduction in HSV1 attachment to HS. Most importantly, such an inhibition process did not dominate in our multiple cycle infection experiment, given the pronounced acceleration if HSV1 growth reported in Fig. 1B and the qualitatively similar proviral effect observed in the single (Fig. 1C) and multiple (Fig. 1B) steps infection experiment. It is also worth to consider that HSV1 can employ both cell-free and cell-to-cell transmission (Dingwell et al., 1994, 1995; Howard et al., 2014). In the later case, HSV1 transmission bypasses the initial attachment to the cell surface, the step hindered by HS-bound ApoE.

Our observation that coating of HSV1 with ApoE leads to enhanced attachment to the cell surface suggests that viruses produced from an ApoE-expressing cell may exhibit enhanced cell-free transmission to target cells, regardless of the later express ApoE or not. This may be of particular clinical relevance, considering that HSV1 primarily infects epidermal keratinocytes where ApoE is naturally expressed and secreted (Fenjves et al., 1989), followed by the infection of sensory neurons and retrograde transportation along the axons to the cell body of neurons (Antinone & Smith, 2010) where ApoE is generally lacking.

In our study, we observed an isoform-dependent pro-viral effect, with ApoE 3 and 4 demonstrating more efficient pro-viral effects than ApoE2 (Fig 1), which is not the case for HCV (Cun et al., 2010). These isoform-dependent effects correlate with the binding affinities of the different ApoE isoforms to heparin, which originally inspired us to assume that virus-bound ApoE manipulates HSV1-HS interactions. Indeed, the binding affinities of ApoE to heparin vary among different isoforms: ApoE 4 and 3 have similar affinities which are higher than ApoE 2 (Futamura et al., 2005). In addition to this, we report that ApoE 4 associates more firmly than ApoE 2 and 3 to the HSV1 virion (Fig. S5), an observation which may further contribute to explain the increased proviral effect of ApoE 4.

ApoE is an important host protein which has been shown to modify the infection process of a variety of viruses. Our results reveal novel roles of ApoE during HSV1 infection at molecular levels, and some isoform dependent effects, which can be likely attributed to the binding abilities of different ApoE isoforms to HS. Although interactions between ApoE 4 and HSV1 have been correlated to increased risk of AD development, the interactions between ApoE and HS have not been linked to the disease. Thus, the significance of ApoE and HSV1 interactions in AD development remains to be studied, considering that AD is a complicated process of little known aetiology. Especially, other factors such as amyloid-beta and tau protein should be considered when studying the development of the disease. In addition, ApoE is naturally expressed and secreted by epidermal keratinocytes (Fenjves et al., 1989), which indicates that the proviral effects of ApoE presented here may also be of relevance during primary and/or recurrent infection in the epithelium. Considering the limitations of data generated from immortalized cell culture, more clinically relevant and comprehensive models, such as epidermal keratinocytes and recently developed brain organoids (Bi et al., 2021; Martens et al., 2021), are suggested for further and deeper studies of ApoE and HSV1 interactions, as well as their significance in disease development.

## Materials and method

### Cells, ApoE proteins, glycosaminoglycan, and other proteins

Green monkey kidney (GMK AH-1) cells were kindly provided by Tomas Bergström (Gothenburg University), which has originally described previously (GUENALP, 1965). Neuroblastoma SH-SY5Y (CRL-2266) cells were purchased from ATCC, and kindly provided by Niklas Arnberg (Umeå University). The hepatocyte-derived carcinoma cell line (Huh-7) was a gift from Magnus Evander (Umeå University). GMK and Huh-7 cells were cultured in DMEM (D5648-10L, Sigma), supplemented with 10% fetal bovine serum (FBS, SV30160.03, cytiva), 20 mM HEPES, penicillin (0.5 unit/mL) and streptomycin (50 µg/mL) (P0718, Gibco). SH-SY5Y cells were cultured in a mixture of medium with 1:1 DMEM + F-12 HAM (21700-075, Gibco), 10% FBS, 20 mM HEPES, penicillin (0.5 unit/mL) and streptomycin (50 µg/mL). The ApoE 4 inducible cell line was established by using the Flp-In ^TM^ T-Rex ^TM^ system (a kind gift from Anna Överby group, Umeå university), according to the manufacturer’s instructions (Invitrogen). In brief, the ApoE 4 gene was synthesized and verified by the Protein Expression Platform at Umeå university, and then constructed in the backbone of pcDNA™5/FRT/TO expression vector. Together with a recombinase expressing vector pOG44, pcDNA™5/FRT/TO-ApoE 4 was co-transfected into a commercially established Flp-In™-293 cell line (based on HEK-293) carrying the integrated FRT. After transfection, selection for successful gene integration was done under hygromycin (100 µg/mL). The ApoE 4 inducible cell line was derived from a single clone and the induction of ApoE 4 expression by tetracycline (1 µg/mL) was verified by western blot. Cell cultures were maintained 37°C and 5% CO_2_ in an incubator. Lyophilized recombinant ApoE 2 (AE-100-10), 3 (AE-101-10), and 4 (AE-102-10) were obtained from AlexoTech AB (Umeå, Sweden). To prepare the protein solution, lyophilised ApoE proteins was first dissolved in 20 mM NaOH and then mixed with 10× PBS (phosphate buffer saline; Medicaco AB, Sweden) (volume ratio of 20 mM NaOH and 10× PBS is 9:1)) to make the stock solution at a concentration of 50 µM (= 1.7 mg/mL). ApoE protein solutions were aliquoted and kept at −80°C. The dilution buffer alone was used in control experiments, termed as the diluent group. All protein aliquots used for experiments experienced no more than two freeze-thaw cycles, to minimise variations in protein concentrations.

Lyophilized parental HS (GAG-HS01 BN1, Iduron, UK) with an average molecular weight of 40 kDa was gently mixed in milli-Q water (Millipore integral system, Molsheim, France) overnight at 4°C to produce the stock concentration of 25 µg/mL. Later, the HS sample was biotinylated at its reducing end by oxime oxidation reaction as described in (Thakar et al., 2014) and stored at −20°C until use.

Lyophilized streptavidin (Sigma) was dissolved in milli-Q water at 5 mg/mL, aliquoted and stored at −80°C. Thawed aliquots was used within a week and kept at 4°C till use.

### Viruses and virus purification

Herpes simplex virus 1 (HSV1) strain KOS (VR-1493; ATCC, Manassas, VA) was produced and purified in our lab by ultracentrifugation through sucrose gradients (Marconi & Manservigi, 2014). In brief, propagation started with MOI 0.01 infection of GMK cells. Viruses from both cells and medium were collected, concentrated, and loaded on sucrose gradients (consist of 2 mL each of 50%, 40%, 30% sucrose layers, w/v) for purification. Cell-associated viruses were released by three freeze-thaw cycles. After centrifugation at 20,200 rpm for 2 h, virus particles sediment at the interface between the 50% and 40% sucrose layers. After collection of the virus-containing fraction, viruses were aliquoted in sucrose for infection studies on cultured cells. For biophysical experiments, sucrose was exchanged with PBS by an extra round of high-speed centrifugation (48,384 g, 20 min at 4°C).

### Quantification of cell-associated virus

To quantify the total amount of cell-associated viruses, infected cells were harvested separately from the medium for DNA extraction and subsequent qPCR quantification of viral genome copies. DNA extraction was done according to the manufacturer’s instruction (Invisob Spin Virus DNA Mini Kit, IBL). Virus DNA was quantified by qPCR with primers targeting the US5 (unique short 5) (Filén et al., 2004) gene as previously described (Abidine et al., 2022). A standard curve established with pI18-HSV1-US5 as the template was used to calculate the absolute quantities of HSV1 copies. The two primers used for qPCR were: HSV1-US5-F, (5’-GGCCTGGCTATCCGGAGA-3’); HSV1-US5-R (5’-GCGCAGAGACATCGCGA-3’). The probe for qPCR was 5’-6FAM-CAGCACACGACTTGGCGTTCTGTGT-Dark Quencher-3’. The qPCR program run as 95°C, 3 min; 40 cycles (95°C, 15 s; 60°C, 30 s). To specifically quantify virus particles on the cell surface, viruses were released by trypsin digestion and separated from cells by centrifugation. Experiments were carried out in 12-well plates. At indicated time points after infection, medium samples were harvested for qPCR or titration by plaque assay. Cells were then washed with PBS twice, followed by trypsin addition. After trypsin digestion, cells were resuspended in PBS and collected for centrifugation at 1,500 rpm for 5 min. After centrifugation, cells and supernatants were separated. Released viruses in supernatants and intracellular viruses were then processed for DNA extraction and genome quantification by qPCR.

### Quantification of virus release rate from infected cells

The release rate of virus was investigated in the premise that similar amounts of viruses were found on the cell surface (Fig. 2B) in all groups at the indicated starting time point. The virus release into the medium (Fig. 2C, without refreshing of any ApoE) was then monitored for 1 or 2 h, after the removal of the supernatants and 3 times wash of cells with ice-cold PBS. Released viruses were later quantified by plaque assay.

### Co-sedimentation, SDS-page, and western blot

Co-sedimentation analysis of HSV1 and ApoE was done in a similar procedure as for the virus production and purification described above. Medium of HSV1-infected Huh-7 cells (only released HSV1) or mock was collected and concentrated, followed by separation by ultracentrifugation through sucrose gradients (consisting of 2 mL each of 55%, 50%, 45%, 40%, 35% sucrose layers, w/v). After centrifugation, samples loaded on top of the sucrose gradients were harvested as one fraction, termed as “solution (S)”. Samples in the sucrose gradients were fractioned from top to bottom (1 mL/fraction).

Samples from each fraction were processed for SDS-page and western blot for detection of ApoE (PA5-27088, Invitrogen, diluted in 5% milk in PBS-tween) and HSV1-gC (B1C1B4, diluted in 5% milk in PBS-tween) (Delguste et al., 2019). In brief, samples for SDS-page and western blot were lysed in lysis buffer (0.05 M Tris-HCl pH=8, 0.15 M NaCl, 1% Triton X-100, in H_2_O + protease inhibitor cocktails (Roche)) for 15 min on ice, followed by centrifugation at 12,000 rpm for 15 min, at 4°C. The supernatants were collected, and aliquots were mixed with 6x laemmli buffer then boiled at 95°C for 10 min. Boiled samples were quickly cooled down on ice and used for SDS-page. Intensities of ApoE and HSV1-gC bands were analysed by ImageJ (Schindelin et al., 2012). Titres of the corresponding fractions were quantified by plaque assay for their infectivity.

### Purified HSV1 and ApoE 4 incubation, complex separation, and fluorescence labelling

Incubation and fluorescence labelling of purified HSV1 and ApoE 4 protein was done at the same time for the experiments described in Fig. 3, 5 and 6. 2×10^8^ PFU HSV1, 8µL ApoE (50 µM in stock), and 1 µL fluorescent lipophilic dye 1,1’-dioctadecyl-6,6’-di(4-sulfophenyl)-3,3, 3’,3’-tetramethylindocarbocyanine (SP-DiIC18(3), or in short SP-DiI) (D7777, Invitrogen, 400 µM in stock) were mixed in PBS in a 1.5 mL Eppendorf tube with the final volume as 100 µL. The mixtures were incubated under rotation for 1 h at room temperature, followed by adding 100 µL/sample Capto-core beads (Capto Core 700, 17548101, Cytiva) to the same tube and a further 1h incubation at 4°C. Capto-core beads were separated by centrifugation at 800 g, 2 min. The supernatants were collected and further purified by running through S-200 columns (MicroSpin^TM^S-200 HR Columns, 27512001, Cytiva). In parallel, HSV1 mixed with diluent (without ApoE 4 added), was included as a control for comparison in the designated experiments. Labelled HSV1 with ApoE or diluent was kept on ice for a few hours (3 - 4h) before adding to SLBs for experiments. When HSV1 without ApoE incubation was used for kinetic analysis (Fig. 7), 2×10^8^ PFU virus was freshly labelled prior to each experiment with 1 µL of fluorescent lipophilic dye 3,3’-Dioctadecyl-5,5’-Di(4-Sulfophenyl) Oxacarbocyanine, Sodium Salt (SP-DiOC18 (3), in short SP-DiO) (D7778, Invitrogen, 400 µM in stock), followed by size exclusion and buffer exchange by filtration through MicroSpin S-200 HR columns (27-5120-01, Cytiva,) (N. Peerboom, et al., 2017, Biophs. J.; N. Peerboom, et al., 2018, ACS Infect. Dis). After labelling, the amount of lost viruses were estimated to 60% using a Förster Resonance Energy Transfer (FRET)-based assay (Thorsteinsson et al., 2020).

### Virus binding kinetics and efficiency

Virus binding quantification of HSV1 and ApoE 4-coated HSV1 (Fig. 4) was done in 12-well plates by qPCR. To get the same input, freshly prepared (as described above, no freeze-thaw cycles) HSV1 and ApoE 4-coatd HSV1 were quantified by qPCR after DNA extraction (as described above). After quantification, 50,000 copies of each were added to cells for binding synchronization on ice while rocking the plate every 10 min. After the selected periods of binding time (t = 15, 30, 45, 60, 90, and 120 min) on ice, unbound viruses in the solutions were removed and cells were quickly washed with ice-cold PBS twice. The attached viruses were harvested together with cells for DNA extraction and quantification by qPCR.

### Viral entry kinetic and efficiency assay

Virus entry efficiencies in Fig. 4B were investigated by a previously described method (Abidine et al., 2022). Before the entry assay, virus binding was done in the same way as in binding quantification. Based on our previous titration results of HSV1 and ApoE 4-coated HSV1, the inputs were estimated as 200 PFUs for both forms of HSV1. Thereafter, viruses attached to the cell surface were either inactivated immediately (t=0 min) by low pH buffer (pH=3, 40 mM citric acid, 10 mM KCl, 135 mM NaCl) (Desai et al., 1988; Nicola & Straus, 2004) or shifted to 37°C for active entry for selected periods of times (t=15, 30, 60,90 and 120 min) before low pH buffer inactivation. Low pH inactivation was done by adding 350 µL/well of low pH buffer for 2 min on ice, shaking every 20 – 30 s, followed by two PBS washes. After inactivation of the uninternalized virus, cells were covered by 1% agarose in DMEM (5% FBS) for another 3 days culture until obvious plaque formations. A low MOI was chosen to ensure a clear readout of plaque formation. Cells were then fixed with 4% formaldehyde (in PBS) and stained with crystal violet (1% crystal violet in 20% ethanol solution). The number of plaques was then counted for each condition.

When comparing the entry efficiencies of HSV1 and ApoE 4-coated HSV1, the different binding efficiencies of the two virus forms (Fig. 4A) were considered by quantifying the actual amounts of attached viral particles from a parallel group to the experimental groups after 1h synchronization on ice via qPCR. From three independent experimental repeats, the ratios of (HSV1+ApoE 4) /HSV1 were measured as 0.83, 0.59, and 0.75. The normalization of HSV1+ApoE 4 entry was done by dividing the counted plaque numbers by the corresponding ratios (Fig. 4B).

### Small Unilamellar Vesicles for (native) supported lipid bilayers

1-palmitoyl-2-oleoyl-glycero-3-phosphocholine (POPC) (850457P), 1,2-dioleoyl-sn-glycero-3-[(N-(5-amino-1-carboxypentyl) iminodiacetic acid) succinyl] (18:1 DGS-NTA(Ni)) (790528P), 18:1 Biotinyl Cap PE (DOPE-Cap-β) (870273P), and N-palmitoyl-sphingosine-1-{succinyl[methoxy(polyethylene glycol)5000]} (PEG) (880280P) were purchased from Avanti Polar Lipids (Alabaster, AL, USA). 1,2-dihexadecanoyl-sn-glycero-3-3phosphoethanolamine, triethylammonium salt (TxRed) was purchased from ThermoFisher Scientific (T1395MP). Small unilamellar vesicles (SUVs) were prepared by extrusion using a mini extruder equipped with a 1 mL syringe (610020 and 610017; Avanti Polar Lipids) as described previously (Peerboom et al., 2017). Pure POPC vesicles, POPC:PEG vesicles (99.5:0.5, molar ratio), and POPC:TxRed vesicles (99:1, molar ratio) were all prepared in PBS by extrusion through a 100 nm polycarbonate membrane at least 11 times. POPC:TxRed was prepared at a stock concentration of 4 mg/mL, the others at 1 mg/mL. POPC:DGS-NTA vesicles (96:4, molar ratio) and POPC:DOPE-Cap-β (90:10, molar ratio) were prepared in HEPES saline buffer (HBS, 10 mM HEPES and 150 mM NaCl; pH 7.5) and PBS, respectively at a stock concentration of 1 mg/mL by extrusion through a 50 nm polycarbonate membrane at least 21 times and were stored under N_2_. All stocks were stored at 4°C.

### Binding assay of HSV1 to surface immobilised His-tagged ApoE

Borosilicate glass cover slides with a diameter of 22 mm (631-0158P, round, No.1, VWR) were cleaned using 7x detergent (MP Biomedicals, CA) and milli-Q water with 1:6 (volume ratio) solution close to boil for 2 h, rinsed extensively and stored in milli-Q water. Before use, the slides were rinsed with milli-Q water, N_2_ dried, and treated in a UV/ozone (UV Ozone Cleaner -ProCleaner™ Plus, Bioforce, IA, USA) for 30 min.

Supported lipid bilayers (SLBs) were formed from SUVs by the method of vesicles spreading, through 30 min exposure of 50 µg/mL vesicle solution in HBS supplemented with 10 mM NiCl_2_ to freshly cleaned glass cover slip. The cover slip was fixed onto a custom Teflon holder using a bi-component Twinsil glue (Picodent, Germany), creating eight wells of equal volume. All incubation steps were performed in still solution. The excessive SUV material was removed from each well by rinsing 20 times with 100 µL PBS. After rinsing, SLBs were incubated with POPC vesicles for 30 min at a final concentration of 37.5 µg/mL to ensure the formation of a good quality bilayer by filling up any potential gaps. After rinsing in PBS, six wells were exposed to His-tagged ApoE 4 at a final concentration of 100 µg/mL for 30 min. Wells that were not exposed to His-tagged ApoE 4 were used as control. After rinsing with PBS, six wells were exposed for 30 min to fluorescently labelled HSV1 particles with or without preincubation with ApoE 4 (see purified HSV1 and ApoE incubation section for details). Images (704 x 704 pixels with a 0.183 µm pixel width) of bound particles at six randomly selected positions were collected using TIRFM with an inverted Nikon (Japan) Eclipse Ti-E2 microscope, an oil-immersion 60X objective (Nikon, NA: 1.49) and a 561 nm laser. The images were analysed using ImageJ. A Gaussian blur (sigma:1 pixel) was applied to all images to reduce the noise and particles were counted after thresholding. Raw particle number were normalized between samples to account for differences in the stock concentration. Experiments were repeated twice.

### Determination of the relative concentrations of fluorescently labelled HSV1 samples

The concentration of labelled particles in each sample was calculated using an assay referred to as “bouncing particle analysis”. In brief, this method measures the number of particles that transiently diffuse close to the glass surface into the TIRF excitation volume without interacting with the substrate, i.e., “bouncing” off the surface. Since particles randomly diffuse in the solution, the number of particles detected during this procedure is linearly proportional to the particle concentration in solution. The measurement was performed on pure POPC SLBs, which are resistant to HSV1 attachment, to minimise the interaction between the particle and the substrate. The timelapses were recorded in TIRF mode with the 60X oil immersion objective using a 561 nm laser with at 50% power and at 10 fps for 1min. The movies were analysed using an in-house MATLAB (MathWorks, USA) script using the same principle as described for the kinetic analysis (see section Virus binding kinetics). Only particles detected on the surface for less than 10 frames or 1s were considered non-interacting with the substrate and counted as “bouncing particle” events. The cumulative sum of new bouncing particles was then calculated versus time and fit to determine the rate of arrival (slope of the cumulative sum). This factor is linearly proportional to the particle concentration in solution and was used to normalize the concentrations across samples.

For each sample, movies were recorded at three different randomly selected positions, analysed, and averaged to extract the arrival rate.

### Membrane extraction and purification

Native membrane vesicles (NMVs) were prepared according to a protocol adapted from Pace et al. (Pace et al., 2015). Confluent cells were rinsed with cold PBS, and then harvested on ice in ice-cold harvest buffer (PBS mixed with EDTA-free protease inhibitor cocktail, Roche) using a cell scraper. Harvested cells were pelleted via centrifugation at 600 g for 10 min, before they were disrupted by 4 passes through a CF1 continuous flow cell disruptor (Constant Systems, U.K.) set at 2,500 psi. The suspension was then centrifuged at 2,000 g for 10 min to pellet nuclei and larger organelles, and then at 6,000 g for 20 min to pellet mitochondria. Finally, the suspension was centrifuged at 150,000 g for 90 min to pellet the NMVs.

NMV pellets were resuspended in harvest buffer and purified via sucrose gradient, to separate plasma membrane vesicles from organelle membrane vesicles. The suspension was mixed with equal volume of 80% sucrose solution, and then layered with a 30% and 5% sucrose solution respectively. All sucrose solutions were made with harvest buffer. The sucrose cushion was then centrifuged at 273,000 g for 2.5 h. The NMVs form a hazy band at the 5% and 30% sucrose interface, which was harvested, flash-frozen, and stored at −80°C. All centrifugations were done at 4°C.

Concentration of NMV material was measured using a FRET based assay described previously (Thorsteinsson et al., 2020).

### Native supported bilayer preparation

Native vesicles were mixed with synthetic POPC:PEG vesicles in a 15:85 % ratio, according to the surface area determined with the FRET assay (Thorsteinsson et al., 2020). Mixing was done in 1.5 mL microcentrifuge tubes to a final concentration of 200 µg/mL and a final volume of 60 µL. The mixture was sonicated for 20 min at 40°C in a bath sonicator (37 kHz, Elmasonic S40H, Germany). After sonication, vesicles were then mixed with 0.3 µL of 0.5 µg/mL POPC:TxRed vesicles.

Borosilicate glass cover slides with diameter of 24 mm (631-0161, round, No.1, VWR) were cleaned using 7x as described above, but without UV-ozone treatment. Bilayers were formed in custom-made 10 µL PDMS wells attached to the cover slides. 10 µL of vesicle mixture was injected into the wells and incubated for 30 min at 37°C. Afterwards, wells were rinsed 10x with 10 µL PBS, before passivating the bilayer with POPC vesicles at a final concentration of 50 µg/mL. Bilayers were then visually inspected for unruptured POPC:TxRed vesicles to confirm bilayer formation before proceeding.

### Surface immobilization of heparan sulfate

For TIRF microscopy-based kinetic experiments with surface-bound HS films, round glass cover slips were cleaned, UV treated and glued on Teflon holder as described in the section on the binding assay of HSV1 to surface immobilised His-tagged ApoE SLBs containing 10% biotinylated lipids (POPC:DOPE-Cap-β (90:10, molar ratio)) were formed at final concentration of 100 µg/mL. After SLB formation, each well was rinsed 14x with 150 µL PBS and then incubated with pure POPC vesicles at 50 µg/mL for 30 mins to get a good quality SLBs. After PBS rinse, each well was incubated with streptavidin in PBS for 30 mins at a final concentration of 50 µg/mL. After rinsing, each well was incubated with the final 2% v/w paraformaldehyde (PFA, VWR) for 15 mins to crosslink and prevent lateral diffusion of streptavidin on the SLBs. Following PBS rinse, each well was incubated with a final concentration of 10 µg/mL biotinylated HS (b-HS) for 30 mins. After rinsing, all wells were incubated with 50 µg/mL pure POPC vesicles for ∼1 h before incubating with virus samples. Wells were kept in this solution and to further reduce non-specific binding, virus samples were diluted in the same solution (i.e., 50 µg/mL pure POPC vesicles in PBS).

### Virus binding kinetics

Kinetic timelapses were imaged with the Nikon Ti-E2 microscope in TIRF mode using a 60X oil immersion objective and a 561 nm laser. nSLBs or HS films were incubated with fluorescently labeled virus particles for 1 h before recording kinetics at either 15 seconds/frame for 20 mins for 80 frames (data in Fig. 5B), 90 mins for 361 frames (data in Fig. 5C), 1 second/frame for 20 mins for 1201 frames (data in Fig. 6) or 30 seconds/frame for 180 mins for 361 frames (data in Fig. 7). Timelapse field-of-view is 704 x 704 pixels with a resolution of 0.183 µm/pixel.

Movies were stabilised using an in-house MATLAB script. The MATLAB script takes as reference image the first frame and performs an image registration based on maximising the cross correlation between the current and reference frame. The reference frame is updated after one third of the length of the movie to improve the registration of rapidly changing videos. In the case a shift larger than 15 pixels in any direction is detected, the script updates the reference frame to the last analysed frame and repeats the registration. If no improvement is observed, a more computationally intensive multimodal registration algorithm is attempted. In the case of low density of particles, the script average over all the frames and crops the field of view to only consider the area presenting the 10 brightest spots to reduce possible errors in the correlation calculation due to random noise. Only rigid translation is considered in every case.

We then performed a kinetic analysis of the stabilised movies using an in-house MATLAB script with improved detection capabilities from what previously described (Peerboom et al., 2017). In brief, in each frame, particles are detected by performing two successive 2D peak scan. Firstly, peaks of prominence higher than a threshold set by the user are identified as particles. The threshold is then lowered by a percentage set by the user and newly detected peaks are considered only if a particle was also detected in the proximity in the previous frame. This is to account for possible bleaching in long exposure experiment which might reduce the intensity of particle on the surface below the higher threshold. The same thresholds were used for all samples for the same experiments. A nearest neighbour algorithm is then used for linking the particle by comparing successive frames with a maximum linking distance of 2 pixels. Possible errors in linking causing the fragmentation of single tracks, thus false attachment, and detachment events, are corrected by comparing the position of tracks ending and beginning in successive frames and linking them if within 1 pixel. Finally, detachment events causing an intensity drop lower than three times the standard deviation of the background are discarded.

For the kinetic analysis, a particle was considered bound if it remained on the surface for at least three consecutive frames. The particle arrival rate was determined by fitting the plot displaying the number of newly detected particles over time with a linear fit (y = Ax + B). To ignore artifacts in the analysis of the initial binding events, the first 13% of data points are excluded from the fitting. The particle dissociation rate was quantified by plotting the number of bound particles as a function of the residence time on the surface and fitting the curve with a double exponential function decay with an offset (*f*(*t*) = A_1_ · exp(−*k*_off_^1^ · *t*) + A_2_ · exp(−*k*_off_^2^ · *t*) + *y*_0_), where A_1_ and A_2_ = amplitude and *y*_0_ = offset to extract two dissociation rates and irreversible fraction. The irreversible fraction was calculated using the following expression: 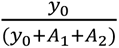. Only particles that bind during the first half of the timelapse were included in the analysis, to avoid biasing the data towards shorter residence times. All binding experiments were normalized to a control which was the binding of naked HSV1 to nSLBs (Fig. 5), to HS films (Fig. 6), or to nSLBs from HEK cells without the induction of ApoE 4 (HEK-) (Fig. 7). Each data point was normalized to its respective control from the same experiment to account for day-to-day variability. After normalization, the results were presented as relative values for association and dissociation rates, and irreversible fraction.

## Acknowledgements

The authors would like to thank Tomas Bergström (Gothenburg university), Sigvard Olofsson (Gothenburg university), Jan Olsson (Umeå University) and Fredrik Elgh (Umeå University) for providing advice during the project development. Magnus Evander, Niklas Arnberg, and Anna Överby (Umeå University) are acknowledged for providing cells and plasmids. We also thank the Biochemical Imaging Center at Umeå University and the National Microscopy Infrastructure, NMI (VR-RFI 2019-00217) for providing assistance in microscopy. This project has been funded by the the Kempe foundation, the Knut and Alice Wallenberg foundation, Åhlensfonden, Alzheimerfonden, Norrländska hjärtfonden, Söderbergs stiftelse (M55/22), and the Swedish research council (2017-04029; 2020-06242).

## References

Abidine, Y., Liu, L., Wallén, O., Trybala, E., Olofsson, S., Bergström, T., & Bally, M. (2022). Cellular Chondroitin Sulfate and the Mucin-like Domain of Viral Glycoprotein C Promote Diffusion of Herpes Simplex Virus 1 While Heparan Sulfate Restricts Mobility. Viruses, 14(8), 1836. 10.3390/v14081836

Ahmad, I., & Wilson, D. W. (2020). Hsv-1 cytoplasmic envelopment and egress. International Journal of Molecular Sciences, 21(17), 1–34. 10.3390/ijms21175969

Altgärde, N., Eriksson, C., Peerboom, N., Phan-Xuan, T., Moeller, S., Schnabelrauch, M., Svedhem, S., Trybala, E., Bergström, T., & Bally, M. (2015). Mucin-like region of herpes simplex virus type 1 attachment protein glycoprotein C (gC) modulates the virus-glycosaminoglycan interaction. Journal of Biological Chemistry, 290(35), 21473–21485. 10.1074/jbc.M115.637363

Antinone, S. E., & Smith, G. A. (2010). Retrograde Axon Transport of Herpes Simplex Virus and Pseudorabies Virus: a Live-Cell Comparative Analysis. Journal of Virology, 84(3), 1504–1512. 10.1128/jvi.02029-09

Ball, M. J. (1982). “Limbic predilection in Alzheimer dementia: is reactivated herpesvirus involved?”. The Canadian Journal of Neurological Sciences. Le Journal Canadien Des Sciences Neurologiques, 9(3), 303–306. 10.1017/s0317167100044115

Bally, M., Gunnarsson, A., Svensson, L., Larson, G., Zhdanov, V. P., & Höök, F. (2011). Interaction of Single Viruslike Particles with Vesicles Containing Glycosphingolipids. Phys. Rev. Lett., 107(18), 188103. 10.1103/PhysRevLett.107.188103

Banfield, B. W., Leduc, Y., Esford, L., Visalli, R. J., Brandt, C. R., & Tufaro, F. (1995). Evidence for an Interaction of Herpes Simplex Virus with Chondroitin Sulfate Proteoglycans during Infection. Virology, 208(2), 531–539. 10.1006/viro.1995.1184

Bankwitz, D., Doepke, M., Hueging, K., Weller, R., Bruening, J., Behrendt, P., Lee, J. Y., Vondran, F. W. R., Manns, M. P., Bartenschlager, R., & Pietschmann, T. (2017). Maturation of secreted HCV particles by incorporation of secreted ApoE protects from antibodies by enhancing infectivity. J Hepatol, 67(3), 480–489. 10.1016/j.jhep.2017.04.010

Benga, W. J., Krieger, S. E., Dimitrova, M., Zeisel, M. B., Parnot, M., Lupberger, J., Hildt, E., Luo, G., McLauchlan, J., Baumert, T. F., & Schuster, C. (2010). Apolipoprotein E interacts with hepatitis C virus nonstructural protein 5A and determines assembly of infectious particles. Hepatology, 51(1), 43–53. 10.1002/hep.23278

Bhattacharjee, P S, Neumann, D. M., Foster, T. P., Bouhanik, S., Clement, C., Vinay, D., Thompson, H. W., & Hill, J. M. (2008). Effect of human apolipoprotein E genotype on the pathogenesis of experimental ocular HSV-1. Exp Eye Res, 87(2), 122–130. 10.1016/j.exer.2008.05.007

Bhattacharjee, P S, Neumann, D. M., Stark, D., Thompson, H. W., & Hill, J. M. (2006). Apolipoprotein E modulates establishment of HSV-1 latency and survival in a mouse ocular model. Curr Eye Res, 31(9), 703–708. 10.1080/02713680600864600

Bhattacharjee, Partha S., Neumann, D. M., Foster, T. P., Clement, C., Singh, G., Thompson, H. W., Kaufman, H. E., & Hill, J. M. (2008). Effective treatment of ocular HSK with a human apolipoprotein E mimetic peptide in a mouse eye model. Investigative Ophthalmology and Visual Science, 49(10), 4263–4268. 10.1167/iovs.08-2077

Bhattacharjee, Partha S, Neumann, D. M., & Hill, J. M. (2009). A Human Apolipoprotein E Mimetic Peptide Effectively Inhibits HSV-1 TK-Positive and TK-Negative Acute Epithelial Keratitis in Rabbits. Current Eye Research, 34(2), 99–102. 10.1080/02713680802647662

Bi, F. C., Yang, X. H., Cheng, X. Y., Deng, W. Bin, Guo, X. L., Yang, H., Wang, Y., Li, J., & Yao, Y. (2021). Optimization of cerebral organoids: a more qualified model for Alzheimer’s disease research. Translational Neurodegeneration, 10(1), 1–13. 10.1186/s40035-021-00252-3

Boyles, J. K., Notterpek, L. M., & Anderson, L. J. (1990). Accumulation of apolipoproteins in the regenerating and remyelinating mammalian peripheral nerve: Identification of apolipoprotein D, apolipoprotein A-IV, apolipoprotein E, and apolipoprotein A-I. Journal of Biological Chemistry, 265(29), 17805–17815. 10.1016/s0021-9258(18)38235-8

Boyles, J. K., Zoellner, C. D., Anderson, L. J., Kosik, L. M., Pitas, R. E., Weisgraber, K. H., Hui, D. Y., Mahley, R. W., Gebicke-Haerter, P. J., Ignatius, M. J., & Shooter, E. M. (1989). A role for apolipoprotein E, apolipoprotein A-I, and low density lipoprotein receptors in cholesterol transport during regeneration and remyelination of the rat sciatic nerve. Journal of Clinical Investigation, 83(3), 1015–1031. 10.1172/JCI113943

Brown, J. C. (2017). Herpes simplex virus latency: The DNA repair-Centered pathway. Advances in Virology, 2017. 10.1155/2017/7028194

Burgos, J. S., Ramirez, C., Sastre, I., Bullido, M. J., & Valdivieso, F. (2002). Involvement of Apolipoprotein E in the Hematogenous Route of Herpes Simplex Virus Type 1 to the Central Nervous System. Journal of Virology, 76(23), 12394–12398. 10.1128/jvi.76.23.12394-12398.2002

Burgos, J. S., Ramirez, C., Sastre, I., & Valdivieso, F. (2006). Effect of Apolipoprotein E on the Cerebral Load of Latent Herpes Simplex Virus Type 1 DNA. Journal of Virology, 80(11), 5383– 5387. 10.1128/jvi.00006-06

Chang, K. S., Jiang, J., Cai, Z., & Luo, G. (2007). Human apolipoprotein e is required for infectivity and production of hepatitis C virus in cell culture. J Virol, 81(24), 13783–13793. 10.1128/JVI.01091-07

Cun, W., Jiang, J., & Luo, G. (2010). The C-terminal alpha-helix domain of apolipoprotein E is required for interaction with nonstructural protein 5A and assembly of hepatitis C virus. J Virol, 84(21), 11532–11541. 10.1128/JVI.01021-10

Delguste, M., Peerboom, N., Le Brun, G., Trybala, E., Olofsson, S., Bergström, T., Alsteens, D., & Bally, M. (2019). Regulatory Mechanisms of the Mucin-Like Region on Herpes Simplex Virus during Cellular Attachment. ACS Chemical Biology, 14(3), 534–542. 10.1021/acschembio.9b00064

Desai, P. J., Schaffer, P. A., & Minson, A. C. (1988). Excretion of non-infectious virus particles lacking glycoprotein H by a temperature-sensitive mutant of herpes simplex virus type 1: Evidence that gH is essential for virion infectivity. Journal of General Virology, 69(6), 1147– 1156. 10.1099/0022-1317-69-6-1147

Dingwell, K. S., Brunetti, C. R., Hendricks, R. L., Tang, Q., Tang, M., Rainbow, A. J., & Johnson, D. C. (1994). Herpes simplex virus glycoproteins E and I facilitate cell-to-cell spread in vivo and across junctions of cultured cells. Journal of Virology, 68(2), 834–845. 10.1128/jvi.68.2.834-845.1994

Dingwell, K. S., Doering, L. C., & Johnson, D. C. (1995). Glycoproteins E and I facilitate neuron-to-neuron spread of herpes simplex virus. Journal of Virology, 69(11), 7087–7098. 10.1128/jvi.69.11.7087-7098.1995

Dobson, C. B., Sales, S. D., Hoggard, P., Wozniak, M. A., & Crutcher, K. A. (2006). The receptor-binding region of human apolipoprotein E has direct anti-infective activity. Journal of Infectious Diseases, 193(3), 442–450. 10.1086/499280

Dong, L. M., Parkin, S., Trakhanov, S. D., Rupp, B., Simmons, T., Arnold, K. S., Newhouse, Y. M., Innerarity, T. L., & Weisgraber, K. H. (1996). Novel mechanism for defective receptor binding of apolipoprotein E2 in type III hyperlipoproteinemia. Nature Structural Biology, 3(8), 718– 722. 10.1038/nsb0896-718

Dong, L. M., & Weisgraber, K. H. (1996). Human apolipoprotein E4 domain interaction. Arginine 61 and glutamic acid 255 interact to direct the preference for very low density lipoproteins. The Journal of Biological Chemistry, 271(32), 19053–19057. 10.1074/jbc.271.32.19053

Elshourbagy, N. A., McLean, J. W., Mahley, R. W., & Taylor, J. M. (1984). Apolipoprotein E mRNA is relatively abundant in the brain, as well as in the liver and other tissues of marmosets and rats. Federation Proceedings, 43(6), 203–207.

Fenjves, E. S., Gordon, D. A., Pershing, L. K., Williams, D. L., & Taichman, L. B. (1989). Systemic distribution of apolipoprotein E secreted by grafts of epidermal keratinocytes: Implications for epidermal function and gene therapy. Proceedings of the National Academy of Sciences of the United States of America, 86(22), 8803–8807. 10.1073/pnas.86.22.8803

Feyzi, E., Trybala, E., Bergström, T., Lindahl, U., & Spillmann, D. (1997). Structural requirement of heparan sulfate for interaction with herpes simplex virus type 1 virions and isolated glycoprotein C. Journal of Biological Chemistry, 272(40), 24850–24857. 10.1074/jbc.272.40.24850

Filén, F., Strand, A., Allard, A., Blomberg, J., & Herrmann, B. (2004). Duplex real-time polymerase chain reaction assay for detection and quantification of herpes simplex virus type 1 and herpes simplex virus type 2 in genital and cutaneous lesions. Sexually Transmitted Diseases, 31(6), 331–336. 10.1097/00007435-200406000-00002

Fleminger, S., Oliver, D. L., Lovestone, S., Rabe-Hesketh, S., & Giora, A. (2003). Head injury as a risk factor for Alzheimer’s disease: The evidence 10 years on; a partial replication. Journal of Neurology Neurosurgery and Psychiatry, 74(7), 857–862. 10.1136/jnnp.74.7.857

Futamura, M., Dhanasekaran, P., Handa, T., Phillips, M. C., Lund-Katz, S., & Saito, H. (2005). Two-step mechanism of binding of apolipoprotein E to heparin: Implications for the kinetics of apolipoprotein E-heparan sulfate proteoglycan complex formation on cell surfaces. Journal of Biological Chemistry, 280(7), 5414–5422. 10.1074/jbc.M411719200

Gastaminza, P., Dryden, K. A., Boyd, B., Wood, M. R., Law, M., Yeager, M., & Chisari, F. V. (2010). Ultrastructural and Biophysical Characterization of Hepatitis C Virus Particles Produced in Cell Culture. Journal of Virology, 84(21), 10999–11009. 10.1128/jvi.00526-10

Griffiths, G., Pfeiffer, S., Simons, K., & Matlin, K. (1985). Exit of newly synthesized membrane proteins from the trans cisterna of the golgi complex to the plasma membrane. Journal of Cell Biology, 101(3), 949–964. 10.1083/jcb.101.3.949

Guenalp, A. (1965). GROWTH AND CYTOPATHIC EFFECT OF RUBELLA VIRUS IN A LINE OF GREEN MONKEY KIDNEY CELLS. Proceedings of the Society for Experimental Biology and Medicine. Society for Experimental Biology and Medicine (New York, N.Y.), 118, 85–90.

Hadigal, S., Koganti, R., Yadavalli, T., Agelidis, A., Suryawanshi, R., & Shukla, D. (2020). Heparanase-Regulated Syndecan-1 Shedding Facilitates Herpes Simplex Virus 1 Egress. Journal of Virology, 94(6). 10.1128/jvi.01672-19

Hadigal, S. R., Agelidis, A. M., Karasneh, G. A., Antoine, T. E., Yakoub, A. M., Ramani, V. C., Djalilian, A. R., Sanderson, R. D., & Shukla, D. (2015a). Heparanase is a host enzyme required for herpes simplex virus-1 release from cells. Nature Communications, 6. 10.1038/ncomms7985

Hadigal, S. R., Agelidis, A. M., Karasneh, G. A., Antoine, T. E., Yakoub, A. M., Ramani, V. C., Djalilian, A. R., Sanderson, R. D., & Shukla, D. (2015b). Heparanase is a host enzyme required for herpes simplex virus-1 release from cells. Nature Communications, 6, 6985. 10.1038/ncomms7985

Herold, B. C., Visalli, R. J., Susmarski, N., Brandt, C. R., & Spear, P. G. (1994). Glycoprotein C-independent binding of herpes simplex virus to cells requires cell surface heparan sulphate and glycoprotein B. Journal of General Virology, 75(6), 1211–1222. 10.1099/0022-1317-75-6-1211

Herold, B. C., WuDunn, D., Soltys, N., & Spear, P. G. (1991). Glycoprotein C of herpes simplex virus type 1 plays a principal role in the adsorption of virus to cells and in infectivity. Journal of Virology, 65(3), 1090–1098. 10.1128/jvi.65.3.1090-1098.1991

Herz, J., & Bock, H. H. (2002). Lipoprotein receptors in the nervous system. Annual Review of Biochemistry, 71, 405–434. 10.1146/annurev.biochem.71.110601.135342

Hogue, I. B., Bosse, J. B., Hu, J.-R., Thiberge, S. Y., & Enquist, L. W. (2014). Cellular mechanisms of alpha herpesvirus egress: live cell fluorescence microscopy of pseudorabies virus exocytosis. PLoS Pathogens, 10(12), e1004535. 10.1371/journal.ppat.1004535

Hopkins, J., Yadavalli, T., Agelidis, A. M., & Shukla, D. (2018a). Host Enzymes Heparanase and Cathepsin L Promote Herpes Simplex Virus 2 Release from Cells. Journal of Virology, 92(23). 10.1128/jvi.01179-18

Hopkins, J., Yadavalli, T., Agelidis, A. M., & Shukla, D. (2018b). Host Enzymes Heparanase and Cathepsin L Promote Herpes Simplex Virus 2 Release from Cells. Journal of Virology, 92(23). 10.1128/JVI.01179-18

Howard, P. W., Wright, C. C., Howard, T., & Johnson, D. C. (2014). Herpes Simplex Virus gE/gI Extracellular Domains Promote Axonal Transport and Spread from Neurons to Epithelial Cells. Journal of Virology, 88(19), 11178–11186. 10.1128/jvi.01627-14

Huemer, H. P., Menzel, H. J., Potratz, D., Brake, B., Falke, D., Utermann, G., & Dierich, M. P. (1988). Herpes simplex virus binds to human serum lipoprotein. Intervirology, 29(2), 68–76. 10.1159/000150031

Itzhaki, R. F., Dobson, C. B., Lin, W. R., & Wozniak, M. A. (2001). Association of HSV1 and apolipoprotein E-ε4 in Alzheimer’s disease. Journal of NeuroVirology, 7(6), 570–571. 10.1080/135502801753248169

Itzhaki, Ruth F., Lin, W. R., Shang, D., Wilcock, G. K., Faragher, B., & Jamieson, G. A. (1997). Herpes simplex virus type 1 in brain and risk of Alzheimer’s disease. Lancet, 349(9047), 241–244. 10.1016/S0140-6736(96)10149-5

Jiang, J., & Luo, G. (2009). Apolipoprotein E but not B is required for the formation of infectious hepatitis C virus particles. J Virol, 83(24), 12680–12691. 10.1128/JVI.01476-09

Jiang, J., Wu, X., Tang, H., & Luo, G. (2013). Apolipoprotein E mediates attachment of clinical hepatitis C virus to hepatocytes by binding to cell surface heparan sulfate proteoglycan receptors. PLoS One, 8(7), e67982. 10.1371/journal.pone.0067982

Kim, J., Basak, J. M., & Holtzman, D. M. (2009). The Role of Apolipoprotein E in Alzheimer’s Disease. Neuron, 63(3), 287–303. 10.1016/j.neuron.2009.06.026

Krastins, B., Prakash, A., Sarracino, D. A., Nedelkov, D., Niederkofler, E. E., Kiernan, U. A., Nelson, R., Vogelsang, M. S., Vadali, G., Garces, A., Sutton, J. N., Peterman, S., Byram, G., Darbouret, B., Pérusse, J. R., Seidah, N. G., Coulombe, B., Gobom, J., Portelius, E., … Lopez, M. F. (2013). Rapid development of sensitive, high-throughput, quantitative and highly selective mass spectrometric targeted immunoassays for clinically important proteins in human plasma and serum. Clinical Biochemistry, 46(6), 399–410. 10.1016/j.clinbiochem.2012.12.019

Kuhlmann, I., Minihane, A. M., Huebbe, P., Nebel, A., & Rimbach, G. (2010). Apolipoprotein e genotype and hepatitis C, HIV and herpes simplex disease risk: A literature review. Lipids in Health and Disease, 9, 1–14. 10.1186/1476-511X-9-8

Lee, J. Y., Acosta, E. G., Stoeck, I. K., Long, G., Hiet, M. S., Mueller, B., Fackler, O. T., Kallis, S., & Bartenschlager, R. (2014). Apolipoprotein E likely contributes to a maturation step of infectious hepatitis C virus particles and interacts with viral envelope glycoproteins. J Virol, 88(21), 12422–12437. 10.1128/JVI.01660-14

Li, Y., & Luo, G. (2021). Human low-density lipoprotein receptor plays an important role in hepatitis B virus infection. PLoS Pathogens, 17(7), 1–21. 10.1371/journal.ppat.1009722

Libeu, C. P., Lund-Katz, S., Phillips, M. C., Wehrli, S., Hernáiz, M. J., Capila, I., Linhardt, R. J., Raffaï, R. L., Newhouse, Y. M., Zhou, F., & Weisgraber, K. H. (2001). New Insights into the Heparan Sulfate Proteoglycan-binding Activity of Apolipoprotein E. Journal of Biological Chemistry, 276(42), 39138–39144. 10.1074/jbc.M104746200

Lin, W.-R., Shang, D., Wilcock, G. K., & Itzhaki, R. F. (1995). Alzheimer’s disease, herpes simplex virus type 1, cold sores and apolipoprotein E4. Biochemical Society Transactions, 23(4), 594S–594S. 10.1042/bst023594s

Lin, W. R., Shang, D., & Itzhaki, R. F. (1996). Neurotropic viruses and Alzheimer disease: Interaction of herpes simplex type I virus and apolipoprotein E in the etiology of the disease. Molecular and Chemical Neuropathology, 28(1–3), 135–141. 10.1007/BF02815215

Linard, M., Letenneur, L., Garrigue, I., Doize, A., Dartigues, J. F., & Helmer, C. (2020). Interaction between APOE4 and herpes simplex virus type 1 in Alzheimer’s disease. Alzheimers Dement, 16(1), 200–208. 10.1002/alz.12008

Lopatko Lindman, K, Weidung, B., Olsson, J., Josefsson, M., Kok, E., Johansson, A., Eriksson, S., Hallmans, G., Elgh, F., & Lovheim, H. (2019). A genetic signature including apolipoprotein Eepsilon4 potentiates the risk of herpes simplex-associated Alzheimer’s disease. Alzheimers Dement (N Y), 5, 697–704. 10.1016/j.trci.2019.09.014

Lopatko Lindman, Karin, Lockman-Lundgren, J., Weidung, B., Olsson, J., Elgh, F., & Lövheim, H. (2022). Long-term time trends in reactivated herpes simplex infections and treatment in Sweden. BMC Infectious Diseases, 22(1), 547. 10.1186/s12879-022-07525-w

Lövheim, H., Norman, T., Weidung, B., Olsson, J., Josefsson, M., Adolfsson, R., Nyberg, L., & Elgh, F. (2019). Herpes Simplex Virus, APOE ɛ4, and Cognitive Decline in Old Age: Results from the Betula Cohort Study. Journal of Alzheimer’s Disease, 67(1), 211–220. 10.3233/JAD-171162

Mahley, R. W., & Rall Jr., S. C. (2000). Apolipoprotein E: far more than a lipid transport protein. Annu Rev Genomics Hum Genet, 1, 507–537. 10.1146/annurev.genom.1.1.507

Marconi, P., & Manservigi, R. (2014). Herpes simplex virus growth, preparation, and assay. Methods in Molecular Biology (Clifton, N.J.), 1144, 19–29. 10.1007/978-1-4939-0428-0_2

Martens, Y. A., Xu, S., Tait, R., Li, G., Zhao, X. C., Lu, W., Liu, C.-C., Kanekiyo, T., Bu, G., & Zhao, J. (2021). Generation and validation of APOE knockout human iPSC-derived cerebral organoids. STAR Protocols, 2(2), 100571. 10.1016/j.xpro.2021.100571

Martínez-Morillo, E., Hansson, O., Atagi, Y., Bu, G., Minthon, L., Diamandis, E. P., & Nielsen, H. M. (2014). Total apolipoprotein E levels and specific isoform composition in cerebrospinal fluid and plasma from Alzheimer’s disease patients and controls. Acta Neuropathologica, 127(5), 633–643. 10.1007/s00401-014-1266-2

Martínez-Morillo, E., Nielsen, H. M., Batruch, I., Drabovich, A. P., Begcevic, I., Lopez, M. F., Minthon, L., Bu, G., Mattsson, N., Portelius, E., Hansson, O., & Diamandis, E. P. (2014). Assessment of peptide chemical modifications on the development of an accurate and precise multiplex selected reaction monitoring assay for Apolipoprotein e isoforms. Journal of Proteome Research, 13(2), 1077–1087. 10.1021/pr401060x

Miller, R. M., & Federoff, H. J. (2008). Isoform-specific effects of ApoE on HSV immediate early gene expression and establishment of latency. Neurobiol Aging, 29(1), 71–77. 10.1016/j.neurobiolaging.2006.09.006

Mingo, R. M., Han, J., Newcomb, W. W., & Brown, J. C. (2012). Replication of Herpes Simplex Virus: Egress of Progeny Virus at Specialized Cell Membrane Sites. Journal of Virology, 86(13), 7084–7097. 10.1128/jvi.00463-12

Müller, H. W., Gebicke-Härter, P. J., Hangen, D. H., & Shooter, E. M. (1985). A Specific 37,000-Dalton Protein That Accumulates in Regenerating but Not in Nonregenerating Mammalian Nerves Abstract. *Science (New York*, N.Y.), 228(4698), 499–501.

Myklebost, O., & Rogne, S. (1988). A physical map of the apolipoprotein gene cluster on human chromosome 19. Human Genetics, 78(3), 244–247. 10.1007/BF00291670

Nicola, A. V, & Straus, S. E. (2004). Cellular and viral requirements for rapid endocytic entry of herpes simplex virus. J Virol, 78(14), 7508–7517. 10.1128/JVI.78.14.7508-7517.2004

Olofsson, S., Bally, M., Trybala, E., & Bergström, T. (2023). Structure and Role of O-Linked Glycans in Viral Envelope Proteins. Annual Review of Virology, 10(1), 1–22. 10.1146/annurev-virology-111821-121007

Pace, H., Simonsson Nyström, L., Gunnarsson, A., Eck, E., Monson, C., Geschwindner, S., Snijder, A., & Höök, F. (2015). Preserved transmembrane protein mobility in polymer-supported lipid bilayers derived from cell membranes. Analytical Chemistry, 87(18), 9194–9203. 10.1021/acs.analchem.5b01449

Pandey, J. P., Olsson, J., Weidung, B., Kothera, R. T., Johansson, A., Eriksson, S., Hallmans, G., Elgh, F., & Lövheim, H. (2020). An Ig γ Marker Genotype Is a Strong Risk Factor for Alzheimer Disease, Independent of Apolipoprotein E ε4 Genotype. Journal of Immunology (Baltimore, Md. : 1950), 205(5), 1318–1322. 10.4049/jimmunol.2000351

Park, D., Lalli, J., Sedlackova-Slavikova, L., & Rice, S. A. (2015). Functional Comparison of Herpes Simplex Virus 1 (HSV-1) and HSV-2 ICP27 Homologs Reveals a Role for ICP27 in Virion Release. Journal of Virology, 89(5), 2892–2905. 10.1128/jvi.02994-14

Peerboom, N., Block, S., Altgärde, N., Wahlsten, O., Möller, S., Schnabelrauch, M., Trybala, E., Bergström, T., & Bally, M. (2017). Binding Kinetics and Lateral Mobility of HSV-1 on End-Grafted Sulfated Glycosaminoglycans. Biophysical Journal, 113(6), 1223–1234. 10.1016/j.bpj.2017.06.028

Peerboom, N., Schmidt, E., Trybala, E., Block, S., Bergström, T., Pace, H. P., & Bally, M. (2018). Cell Membrane Derived Platform To Study Virus Binding Kinetics and Diffusion with Single Particle Sensitivity. ACS Infectious Diseases, 4(6), 944–953. 10.1021/acsinfecdis.7b00270

Plassman, B. L., Havlik, R. J., Steffens, D. C., Helms, M. J., Newman, T. N., Drosdick, D., Phillips, C., Gau, B. A., Welsh-Bohmer, K. A., Burke, J. R., Guralnik, J. M., & Breitner, J. C. S. (2000). Documented head injury in early adulthood and risk of Alzheimer’s disease and other dementias. Neurology, 55(8), 1158–1166. 10.1212/WNL.55.8.1158

Qiao, L., & Luo, G. G. (2019). Human apolipoprotein E promotes hepatitis B virus infection and production. PLoS Pathog, 15(8), e1007874. 10.1371/journal.ppat.1007874

Rasmussen, K. L., Tybjærg-Hansen, A., Nordestgaard, B. G., & Frikke-Schmidt, R. (2016). Plasma levels of apolipoprotein E and risk of ischemic heart disease in the general population. Atherosclerosis, 246, 63–70. 10.1016/j.atherosclerosis.2015.12.038

Rezeli, M., Zetterberg, H., Blennow, K., Brinkmalm, A., Laurell, T., Hansson, O., & Marko-Varga, G. (2015). Quantification of total apolipoprotein E and its specific isoforms in cerebrospinal fluid and blood in Alzheimer’s disease and other neurodegenerative diseases. EuPA Open Proteomics, 8, 137–143. 10.1016/j.euprot.2015.07.012

Saito, H., Dhanasekaran, P., Nguyen, D., Baldwin, F., Weisgraber, K. H., Wehrli, S., Phillips, M. C., & Lund-Katz, S. (2003). Characterization of the heparin binding sites in human apolipoprotein E. Journal of Biological Chemistry, 278(17), 14782–14787. 10.1074/jbc.M213207200

Schindelin, J., Arganda-Carreras, I., Frise, E., Kaynig, V., Longair, M., Pietzsch, T., Preibisch, S., Rueden, C., Saalfeld, S., Schmid, B., Tinevez, J. Y., White, D. J., Hartenstein, V., Eliceiri, K., Tomancak, P., & Cardona, A. (2012). Fiji: An open-source platform for biological-image analysis. Nature Methods, 9(7), 676–682. 10.1038/nmeth.2019

Shieh, M. T., WuDunn, D., Montgomery, R. I., Esko, J. D., & Spear, P. G. (1992). Cell surface receptors for herpes simplex virus are heparan sulfate proteoglycans. J Cell Biol, 116(5), 1273– 1281. 10.1083/jcb.116.5.1273

Siddiqui, R., Suzu, S., Ueno, M., Nasser, H., Koba, R., Bhuyan, F., Noyori, O., Hamidi, S., Sheng, G., Yasuda-Inoue, M., Hishiki, T., Sukegawa, S., Miyagi, E., Strebel, K., Matsushita, S., Shimotohno, K., & Ariumi, Y. (2018). Apolipoprotein E is an HIV-1-inducible inhibitor of viral production and infectivity in macrophages. PLoS Pathogens, 14(11), e1007372. 10.1371/journal.ppat.1007372

Simon, R., Girod, M., Fonbonne, C., Salvador, A., Clément, Y., Lantéri, P., Amouyel, P., Lambert, J. C., & Lemoine, J. (2012). Total ApoE and ApoE4 isoform assays in an Alzheimer’s disease case-control study by targeted mass spectrometry (n = 669): A pilot assay for methionine-containing proteotypic peptides. Molecular and Cellular Proteomics, 11(11), 1389–1403. 10.1074/mcp.M112.018861

Smith, G. W., Hannan, S. F., Scott, P. J., & Simpson, I. J. (1982). Immune complex-like activity associated with abnormal serum lipoproteins in systemic erythematosus. Clinical and Experimental Immunology, 48(1), 8–16.

Steel, A. J., & Eslick, G. D. (2015). Herpes viruses increase the risk of Alzheimer’s disease: A meta-analysis. Journal of Alzheimer’s Disease, 47(2), 351–364. 10.3233/JAD-140822

Strandberg, T. E., Pitkala, K., Eerola, J., Tilvis, R., & Tienari, P. J. (2005). Interaction of herpesviridae, APOE gene, and education in cognitive impairment. Neurobiology of Aging, 26(7), 1001–1004. 10.1016/j.neurobiolaging.2004.09.008

Thakar, D., Migliorini, E., Coche-Guerente, L., Sadir, R., Lortat-Jacob, H., Boturyn, D., Renaudet, O., Labbe, P., & Richter, R. P. (2014). A quartz crystal microbalance method to study the terminal functionalization of glycosaminoglycans. Chemical Communications, 50(96), 15148–15151. 10.1039/c4cc06905f

Thorsteinsson, K., Olsén, E., Schmidt, E., Pace, H., & Bally, M. (2020). FRET-Based Assay for the Quantification of Extracellular Vesicles and Other Vesicles of Complex Composition. Analytical Chemistry, 92(23), 15336–15343. 10.1021/acs.analchem.0c02271

Trybala, E, Liljeqvist, J. A., Svennerholm, B., & Bergström, T. (2000). Herpes simplex virus types 1 and 2 differ in their interaction with heparan sulfate. J Virol, 74(19), 9106–9114. 10.1128/jvi.74.19.9106-9114.2000

Trybala, Edward, Peerboom, N., Adamiak, B., Krzyzowska, M., Liljeqvist, J.-Å., Bally, M., & Bergström, T. (2021). Herpes Simplex Virus Type 2 Mucin-Like Glycoprotein mgG Promotes Virus Release from the Surface of Infected Cells. Viruses, 13(5), 887. 10.3390/v13050887

Tudorache, I. F., Trusca, V. G., & Gafencu, A. V. (2017). Apolipoprotein E - A Multifunctional Protein with Implications in Various Pathologies as a Result of Its Structural Features. Comput Struct Biotechnol J, 15, 359–365. 10.1016/j.csbj.2017.05.003

Wainberg, M., Luquez, T., Koelle, D. M., Readhead, B., Johnston, C., Darvas, M., & Funk, C. C. (2021). The viral hypothesis: how herpesviruses may contribute to Alzheimer’s disease. Molecular Psychiatry, 26(10), 5476–5480. 10.1038/s41380-021-01138-6

Wang, M., Chen, J., & Turko, I. V. (2012). 15N-labeled full-length apolipoprotein E4 as an internal standard for mass spectrometry quantification of apolipoprotein e isoforms. Analytical Chemistry, 84(19), 8340–8344. 10.1021/ac3018873

Wildsmith, K. R., Han, B., & Bateman, R. J. (2009). Method for the simultaneous quantitation of apolipoprotein E isoforms using tandem mass spectrometry. Analytical Biochemistry, 395(1), 116–118. 10.1016/j.ab.2009.07.049

Xu, Y., Martinez, P., Séron, K., Luo, G., Allain, F., Dubuisson, J., & Belouzard, S. (2015). Characterization of Hepatitis C Virus Interaction with Heparan Sulfate Proteoglycans. Journal of Virology, 89(7), 3846–3858. 10.1128/jvi.03647-14

Yamauchi, Y., Deguchi, N., Takagi, C., Tanaka, M., Dhanasekaran, P., Nakano, M., Handa, T., Phillips, M. C., Lund-Katz, S., & Saito, H. (2008). Role of the N- and C-terminal domains in binding of apolipoprotein E isoforms to heparan sulfate and dermatan sulfate: A surface plasmon resonance study. Biochemistry, 47(25), 6702–6710. 10.1021/bi8003999

Yue, L., & Mazzone, T. (2011). Endogenous adipocyte apolipoprotein E is colocalized with caveolin at the adipocyte plasma membrane. Journal of Lipid Research, 52(3), 489–498. 10.1194/jlr.M011809

