## supporting materials and methods, figures, and table for "Recruitment of apolipoprotein E facilitates Herpes simplex virus 1 attachment, entry, and release"

**[#]Correspondence**

#### Supplementary materials and methods

##### Cell proliferation test

Cell proliferation was evaluated by the WST-1 based colorimetric assay according to the manufacturer's instructions (Roche, 5015944001). In brief, cells were seeded in 96-well plates and cultured overnight. On the second day, medium was changed to 50 µL of new medium containing the designated ApoE and the cells were further cultured for 24 h. 4 h prior to analysis, WST-1 reagent was added (5 µL/well), followed by the measurement of absorbance at 440 nm by an ELISA reader. The corresponding culture medium was included as a control and this readout was subtracted from the experimental groups.

##### Quantification of virus binding

Virus binding quantification (Fig. S4) by qPCR was done in 12-well plates. Prior to adding viruses, cells were treated with ApoE for 4 h. 200 PFUs/well of virus diluted in 150 µL infection medium (DMEM, 1% FBS, 20 mM HEPES, and 1% Penicilline and Streptomycin) was added for infection. Virus binding synchronization was also done on ice while rocking the plate every 10 min for 1 h. The harvest of attached viruses, DNA extraction, and qPCR quantification were described in the materials and methods of the main text. Since the concentration of virus (200 PFUs) was too low for stable detection via qPCR, binding quantification was achieved after increasing the amounts of viruses to 5,000 PFUs and 25,000 PFUs. Quantification of different PFUs as inputs by qPCR revealed an almost perfect linear regression (Abidine et al., 2022), confirming that this approach was viable.

#### Co-immunoprecipitation

Huh-7.5 cells were used for interaction studies between ApoE and HSV1 glycoproteins. Cells in T75 flasks were mock treated or infected with HSV1 at MOI 10. At the indicated times, flasks were placed on ice and washed with ice-cold PBS once. Cells were lysed by adding 1 mL lysis buffer (0.05 M Tris-HCl pH=8, 0.15 M NaCl, 1%Triton X-100, in $H_2O$ + protease inhibitor cocktails (Roche)) to each flask and collected into Eppendorf tubes. The lysis was done by incubating samples on ice for 20 min. Cell debris was separated via centrifugation at 13,000 rpm for 10 min at 4 °C. After centrifugation, 60 µl of cell lysates were kept for input detections according to the method described in the main texts. The rest of the lysates were used for co-immunoprecipitation with anti-ApoE (Invitrogen, PA5-27088) or anti-HSV1 gE antibodies.

Co-immunoprecipitation was done with 30% protein A agarose beads (Sigma, P3476-5), 50 µl/sample. Before incubation, beads were washed 3 times in 1 mL lysis buffer via centrifugation (3,000 rpm, 3 min at 4°C). The washed beads (without adding antibodies) were incubated with cell lysates for 2 h at 4°C to clear unspecific binding. The cleared cell lysates were incubated with 3 µg of ApoE or HSV1-gE antibodies at 4°C overnight. On the following day, new agarose A beads were washed and added to the lysates + antibody solutions for another 90 min incubation at 4°C. Then, the complexes (beads, antibodies, and the attached proteins/protein complexes) were collected and washed with lysis buffer 5 times by centrifugation at 3,000 rpm, 3 min, 4 °C. After the last wash, the lysis buffer was removed with a syringe, and 60 µl 2X Laemmli buffer/sample was added and proceeded for SDS-page and western blot. Antibodies were used to probe ApoE, HSV1-gB (1B11D8), HSV1-gC (B1C1B4), HSV1-gD (C4D5G2), HSV1-gE (B1E6A5) (Delguste et al., 2019), or tubulin (Invitrogen, MA5-16308-HRP).

#### HSV1 binding to immobilised ApoE isoforms

To measure the binding kinetics of HSV1 to surface-immobilised ApoE isoforms, coverslips were cleaned in a boiling solution of 1:10 7x detergent in MilliQ and UV treated. 10 µl PDMS wells were used as described in Material and Methods in the main texts (Native supported bilayer preparation). The SLBs containing 5% NTA-95% POPC were formed by spontaneous rupture of 50 nm vesicles with the same composition at final concentration of 100 µg/mL in HBS supplemented by 10 mM $NiCl_2$. After rinsing, the bilayer was exposed to His-tagged ApoE isoforms (2, 3, and 4) at a concentration of 100 µg/ml for 1 hour. No protein was added for the negative sample. Simultaneously, HSV1 was fluorescently labelled as described in Material and Methods in the main texts. After rinsing the surface in PBS, 5 µl of virus solution was added to 5 µl of PBS remaining in the well and incubated for 1 hour to reach equilibrium. The samples were imaged in TIRFM as described in Material and Methods at 20 s/frame for 1.5 hours. All rinsing steps were performed by adding 10 µl of buffer to a sample volume of 5 µl at least 7 times.

The videos were analysed using the same in-house MATLAB scripts described in Materials and Methods in the main texts. A single exponential fit with offset was used to determine the multivalent dissociation rate constant ($k_{\mathrm{off}}$):

$$y = A\exp(-k_{\mathrm{off}}t) + \mathrm{IF},$$

where IF represent the fraction of particles that behave as irreversibly bond to the substrate in the timescale of the experiment, *A* is a fitting constant and *t* is the elapsed time since particle attachment. The normalised dissociation constant ($\mathrm{K_D}$) was determined by dividing $k_{\mathrm{off}}$ by the association rate for each isoform and by normalising the result by the average value obtained for ApoE 2.

Equal loading of the ApoE isoforms onto the NTA-presenting bilayer and the stability of the resulting bond were confirmed prior to the TIRFM assay using quartz crystal microbalance with dissipation monitoring (QCM-D) (data not shown).

**Viral entry kinetic and efficiency assay**

Virus entry kinetics and efficiency (Fig. S4) were analysed in the same way as described in the materials and methods of the main text. A low MOI (200 PFU) was chosen as the input for all the groups. The normalization of the virus entry was then done by dividing the number of plaques by their corresponding binding factors (binding ratios normalized to the diluent group, Fig. S4a).

### Supplementary figures and table

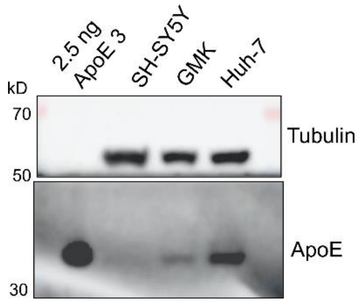


**Fig S1: ApoE expression levels in different cell lines.** Same amounts of SH-SY5Y, GMK, or Huh-7 cells were lysed and analysed by western blot for ApoE expression. Tubulin and ApoE were probed.

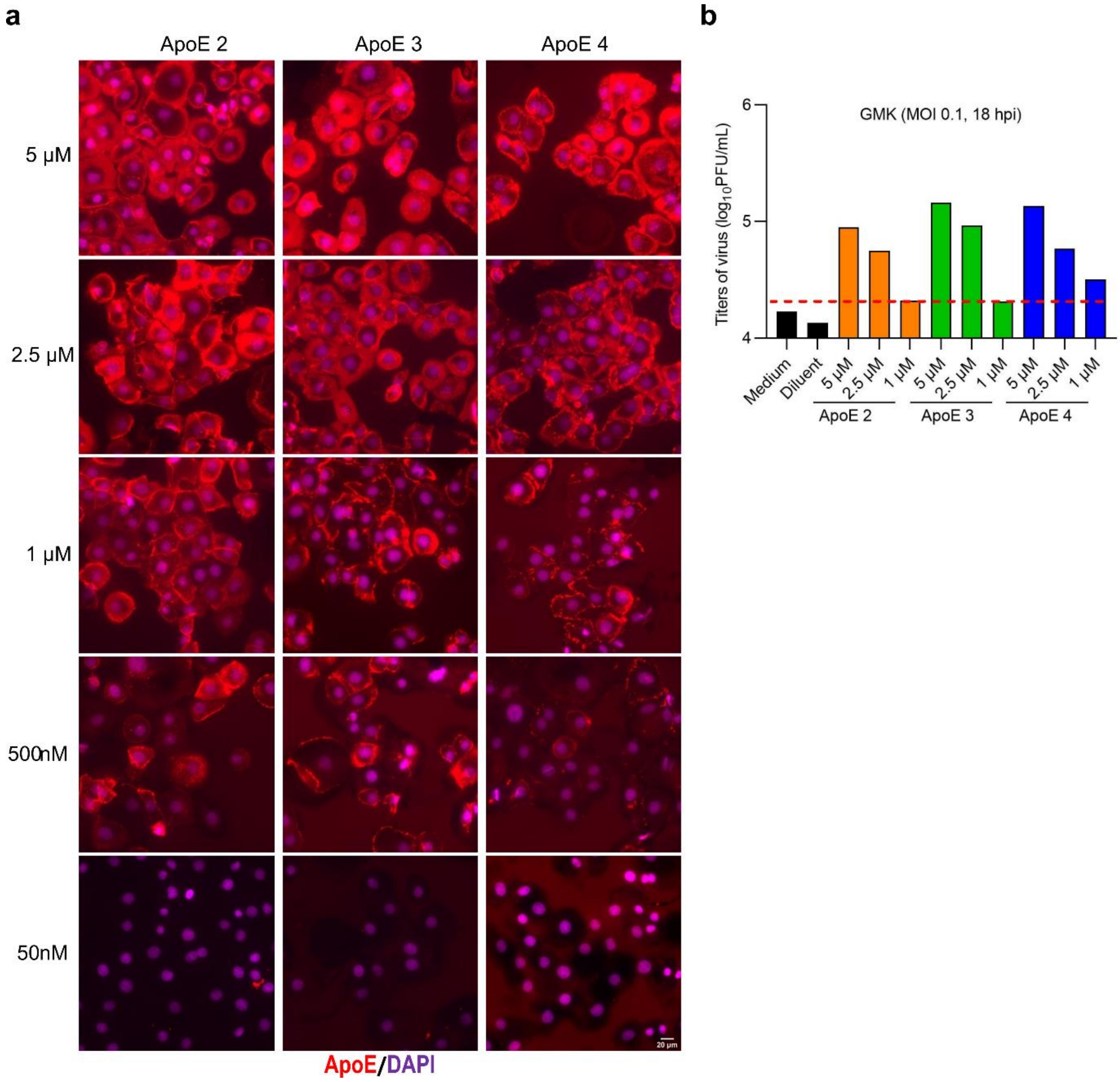

**Fig. S2**: **ApoE uptake and HSV1 infection in GMK cells with different ApoE concentrations (a)** ApoE 2, 3, or 4 was diluted in normal culture medium and added to GMK cells in a 24-well plate with indicated concentrations. After 8 h incubation, cells were fixed with 4% formaldehyde for 15 min and permeabilized with the permeabilization buffer (0.5% Triton X-100, 20 mM glycine in PBS) for 15 min. The fixed samples were incubated with a blocking buffer for 30 min, followed by incubation of primary antibody against ApoE (PA5-27088 ThermoFisher Scientific, 1:1000 in blocking buffer) for 1h and secondary antibody (A32733 ThermoFisher Scientific, Alexa Fluor™ Plus 647, 1:500), together with DAPI staining, for 1h. All the procedures were carried out at room temperature. Images were taken with Nikon (Japan) Eclipse Ti-E2 microscope, 60X oil objective. Representative images were shown for each group. **(b)** HSV1 growth 18 hpi was analysed by plaque assay with indicated concentrations of ApoE 2, 3, or 4 added at 1hpi. Results were generated from a single experiment.

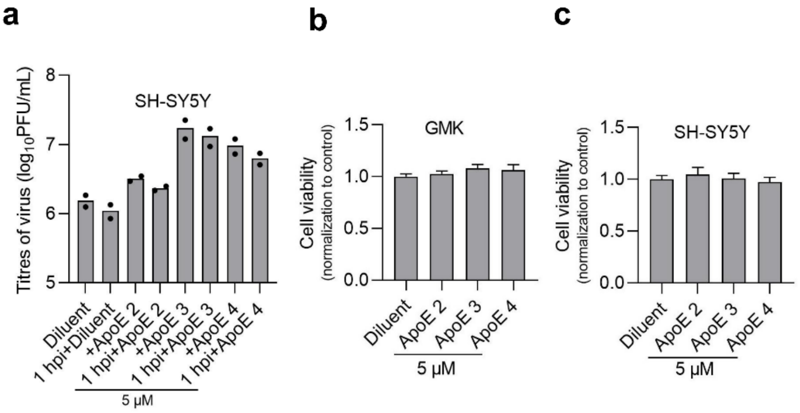


**Fig. S3: HSV1 infection on SH-SY5Y pretreated with ApoE and cell toxicity test after adding ApoE to cells. (a)** HSV1 growth was analysed by plaque assay after infection of SH-SY5Y at MOI 0.1, 24 hpi, where 5 μM ApoE was either pre-incubated overnight (labelled as +ApoE) or added after 1 h virus inoculation (labelled as 1 hpi + ApoE). SH-SY5Y **(b)** or GMK **(c)** cell growth was analysed in the presence of 5 μM ApoE isoforms. The data were normalized to the average absorbance values of the diluent group.

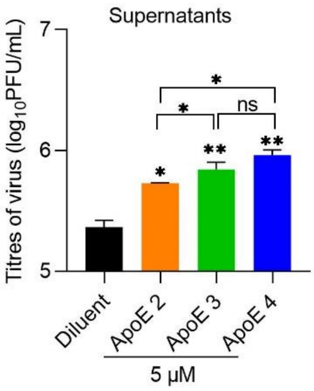


**Fig. S4: HSV1 in the supernatants titrated by plaque assay.** GMK cells were infected with HSV1 (MOI 0.1) and ApoE or diluent was added at 1hpi. At 20 hpi, supernatants and cell samples were separated. Viral genomes in the supernatants (the released HSV1) were quantified by qPCR.

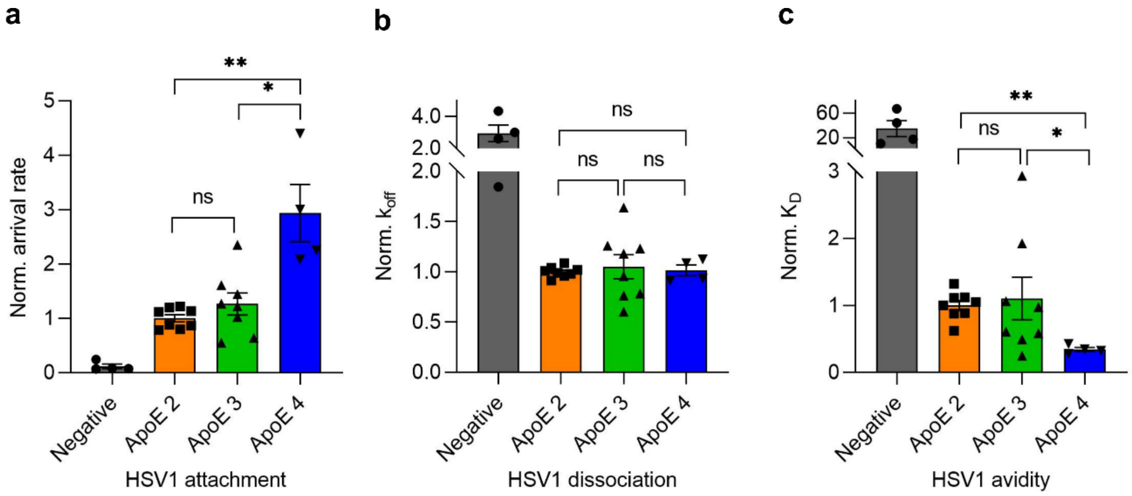


**Fig. S5**: **Kinetic analysis (association rate (a), dissociation rate constant (b) and dissociation constant (c)) of the bond formed between HSV1 and surface-immobilised ApoE isoforms using our TIRFM-based equilibrium fluctuation analysis assay.** ApoE was immobilized on a POPC:DGS-NTA membrane surface as described in conjunction with Fig 3D in the main text. Data were normalised by the average value of ApoE2 for each experimental repeat. Statistical significance determined using Mann-Whitney test between each pair of isoforms. ns: no significance difference detected, *: $p<0.05$, **: $p<0.005$. A substrate lacking ApoE was used as a negative control. The negative control exhibits a significant difference to all other samples in all graphs (not shown to increase readability).

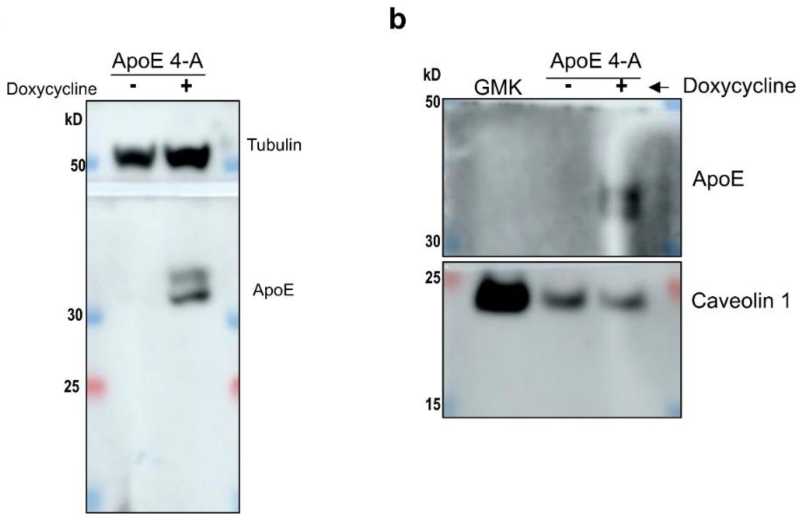


**Fig. S6 Plasma membrane association of ApoE 4 in an ApoE 4 inducible cell line. (a**) Induction of ApoE 4 by doxycycline. ApoE 4 inducible cells were prepared in large culture for membrane extraction. The expression of ApoE 4 was induced by adding doxycycline (1 µg/mL) overnight (about 16 h), before the cells were harvested for membrane extraction. Aliquots of both mock treated and induced cells were collected and analysed by western blot for ApoE expression. Tubulin was probed as a loading control. **(b)** Plasma membrane association of ApoE 4. The presence of ApoE 4 and caveolin 1 were verified by western blot in the extracted and purified plasma membrane samples.

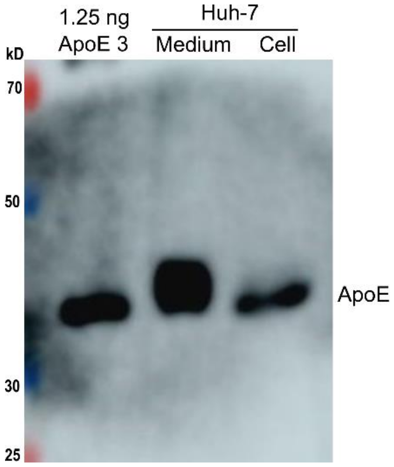


**Fig. S7 ApoE levels in the medium (secreted) and the cells of Huh-7.** Huh-7 cells were cultured in a T75 flask with DMEM (10% FBS, 20 mM HEPES, and penicillin (0.5 unit/mL) and streptomycin (50 µg/mL)). When close to 100% confluence, the culture medium and cells were quantified and collected for lysates preparation and western blot analysis, as described in the methods. The culture medium was 8 mL and cell counts were 3.5 million. 50 µl of medium sample (1/6 was loaded) and 1 million cells (1/48 was loaded) were lysed and prepared for western blot. Purified ApoE 3 (1.25 ng) was included as a control. The concentrations of ApoE in the medium was calculated as 176.574 µg/mL (170 µg/mL = 5 µM); in cells its concentration is 40.43 ng per million cells. Calculations were done based on the intensities of the corresponding bands.

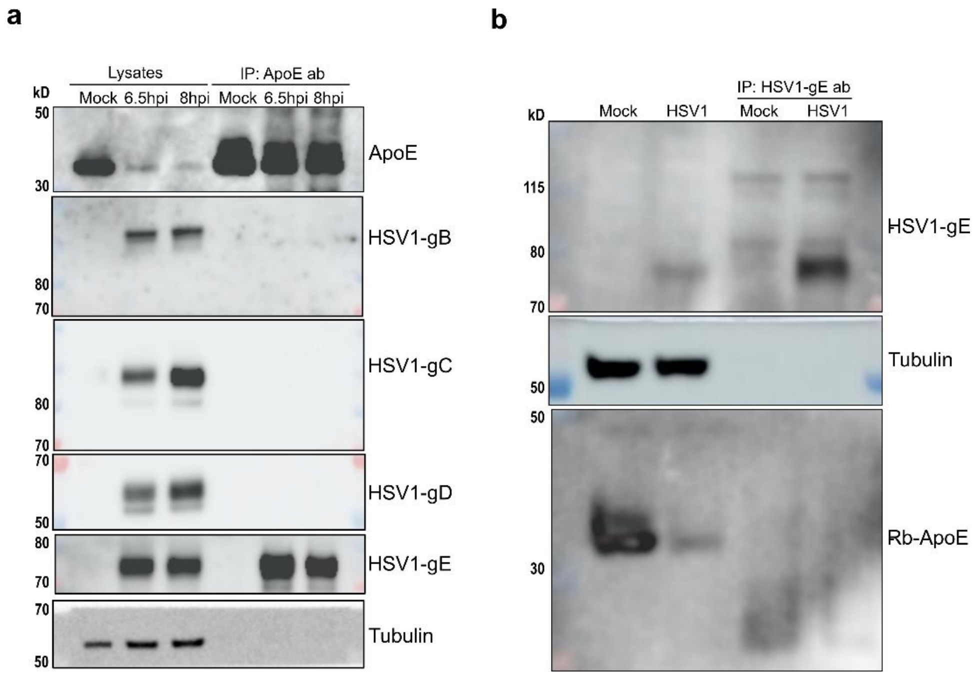


**Fig. S8 Interactions analysis between ApoE and HSV1 glycoproteins.** Huh-7 cells were mock treated or infected with HSV1 (MOI 10). Cells were lysed at indicated time points, followed by immunoprecipitation with antibody against ApoE **(a)** or HSV1-gE **(b)** as described in the methods of the supplementary information. Protein complexes pulled down were separated by SDS page and the indicated proteins were probed by western blot.

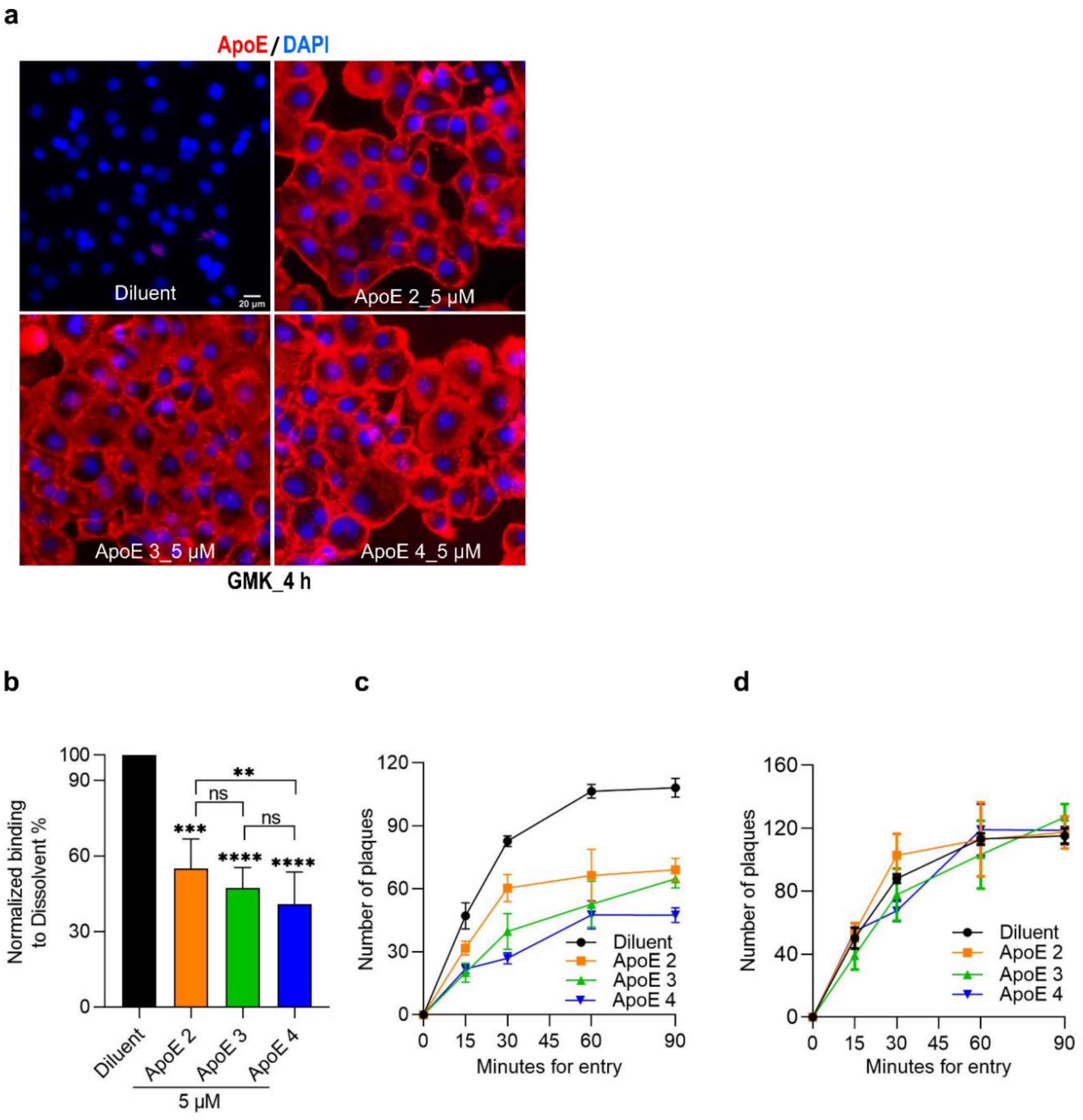


**Fig. S9 (a) ApoE distribution in GMK cells.** Diluent or 5 µM ApoE 2, 3, or 4 was diluted in normal culture medium and added to GMK cells in a 24-well plate. After 4 h incubation, cells were fixed with 4% formaldehyde for 15 min and permeabilized with the permeabilization buffer (0.5% Triton X-100, 20 mM glycine in PBS) for 15 min. The fixed samples were incubated with a blocking buffer for 30 min, followed by incubation of primary antibody against ApoE (PA5-27088 ThermoFisher Scientific, 1:1000 in blocking buffer) for 1 h and secondary antibody (A32733 ThermoFisher Scientific, Alexa Fluor$^{TM}$ Plus 647, 1:500) for 1 h. The nucleus were stained with DAPI together with the secondary antibody incubation. All the procedures were carried out at room temperature. Images were taken with Nikon (Japan) Eclipse Ti-E2 microscope, 60X oil objective. Representative images were shown for each group. **(b, c, and d) HSV1 binding, but not entry, is affected by ApoE when added prior to infection. (b)** HSV1 attached to GMK cells, pre-treated with ApoE or diluent, were quantified by qPCR after 1 h binding synchronization on ice. The data of ApoE groups were normalized to the diluent group. The percentages of ApoE groups are 55% (ApoE 2), 47% (ApoE 3), and 42% (ApoE 4). **(c)** HSV1 entry to GMK treated with ApoE or diluent at selected timepoints were quantified by plaque formation. The average value of the diluent at 90 min was seen as the plateau, to which the rest of the data were normalized. No statistical analysis was compared. **(d)** HSV1 entry efficiencies under different conditions were calculated by dividing the number of plaques by the binding ratios as shown in (a), followed by normalization as done for (b). The normalized entry efficiencies of each ApoE group were compared to the diluent group individually at the same time points. No significance was observed from any of the statistical analysis for (C). Results represent three or more independent repeats. Student t-test, ***: $p \leq 0.001$, and ****: $p \leq 0.0001$. Error bars: mean ± SD.

**Table S1: Summary of the fitted ($A_1$, $A_2$ and $y_0$) and derived ($k_{off}1$ and $k_{off}^2$) parameters** from the double exponential fits of the normalized dissociation curves for the data presented in figures 5, 6 and 7. Each dissociation curve was normalized so that the total number of particles is expressed as a fraction of 1 and then fitted with a double exponential function: $f(t) = A_1 \cdot \exp(-k_{off}^1 \cdot t) + A_2 \cdot \exp(-k_{off}^2 \cdot t) + y_0$). All values are given as mean ± SD (standard deviation) from 3 independent experiments.

| Figure # | Probed surfaces | Fluorescently labelled particles | Fitted parameters | | | Derived parameters | |
|---|---|---|---|---|---|---|---|
| | | | $A_1$ | $A_2$ | $y_0$ | $k_{off}^1$ ($s^{-1}$) | $k_{off}^2$ ($s^{-1}$) |
| 5 | nSLBs | HSV1 | 0.24 ± 0.09 | 0.077 ± 0.040 | 0.841 ± 0.054 | 0.0230 ± 0.0064 | 0.00146 ± 0.00067 |
| | | HSV1+ApoE 4 | 0.40 ± 0.25 | 0.121 ± 0.031 | 0.773 ± 0.082 | 0.0268 ± 0.0057 | 0.00207 ± 0.00075 |
| 6 | HS films | HSV1 | 0.52 ± 0.05 | 0.183 ± 0.078 | 0.620 ± 0.071 | 0.0768 ± 0.0099 | 0.00816 ± 0.0008 |
| | | HSV1+ApoE 4 | 0.64 ± 0.06 | 0.281 ± 0.111 | 0.452 ± 0.133 | 0.0709 ± 0.0217 | 0.00706 ± 0.00099 |
| 7 | HEK-† | HSV1 | 0.47 ± 0.11 | 0.148 ± 0.020 | 0.658 ± 0.063 | 0.0100 ± 0.0016 | 0.00080 ± 0.00014 |
| | HEK+† | | 0.40 ± 0.05 | 0.145 ± 0.027 | 0.708 ± 0.055 | 0.0111 ± 0.0046 | 0.00110 ± 0.00041 |

† HEK- and HEK+ represent the nSLBs from HEK cells without and with the induction of ApoE 4 expression, respectively.
